# Ultra-precision 3D Printed Sample Tubes of Complex Cross Section for Enhanced Electron Paramagnetic Resonance Sensitivity of Aqueous Samples

**DOI:** 10.64898/2026.10.30.763151

**Authors:** Anand Anilkumar, Joseph T. Wherley, Alexander M. Garces, Michael T. Lerch, Candice S. Klug, Jason W. Sidabras

## Abstract

This paper discusses the applications of ultra-precision 3D printing in developing EPR sample tubes for X-band measurements of aqueous samples. The seed idea of this work started from the flat cell idea discussed in Mett and Hyde et al.(2003), Sidabras et al. (2005), and Sidabras et al. (2017), where placing multiple aqueous flat cells perpendicular to the electric field in the microwave cavity reduce RF losses and improve EPR signal sensitivity. In this work, we present the 3D printed aquastar sample holders for dielectric resonators and the aquasun sample holders for cavity resonators. Even though the ultra-precision 3D printing enables the fabrication of complex cross-sections at 25um resolution, the photopolymer material has comparatively higher dielectric losses compared to the conventional materials like glass and PTFE. To reduce the RF losses due to the inherent dielectric properties, we introduced the latticing approach in reducing the amount of plastic without compromising the structural stability. In addition to this, we present the slicing and dielectric approximation approach in reducing the computation cost for these complex cross-sections. The novel sample tubes discussed here present signal improvements through both simulation and experiments, which were validated through power saturation experiments. The sample tube development methodology presented here provide a general framework for designing and implementing 3D printed EPR sample holders, based on resonator electromagnetic field distribution and dielectric properties of the photopolymer material.

## 1. Introduction

Electron Paramagnetic Resonance (EPR) spectroscopy is a prominent technique for studying the structure and dynamics of moieties with a unpaired electron. By utilizing the spin dynamics of this naturally occurring or an engineered unpaired electron, EPR have been widely adopted in structural biology, metalloprotein research, redox research, rational drug design and clinical diagnostics. Specifically for the structural biology studies, the Site Directed Spin Labeling (SDSL) technique, which introduce a spin label to a specific position, which is either a natural cysteine or a genetically engineered cysteine mutation. The protein sample concentration and volume are key considerations while conducting EPR studies. The optimum concentration should be considered as proteins tend to aggregate and several recombinant proteins have very low expression level. Here in this paper, we present the idea of introducing ultra-precision 3D printing in fabricating complex cross-sections relevant for X-band Continuous-Wave (CW) EPR spectroscopy of low concentration protein samples.

The key consideration while placing an aqueous sample in an EPR cavity is to position the sample where the RF electric field is zero, and the RF magnetic field is maximum [1]. Conventionally, X-band EPR measurements of aqueous samples using a Varian commercial rectangular *TE*_102_ cavity employ the thin quartz flat cell cuvettes positioned parallel to the RF electric field at the center of the cavity. Accurate placement of these flat cells are critical, since the misalignment even by few degrees significantly reduce the Q-value. It was reported in Mett et al. that rotating the flat cell precisely by 90^0^ to a perpendicular orientation recovers the Q-value and improves the signal[2]. The rationale for this phenomenon was explained based on three types of losses characterized in Mett & Hyde et al[2]. The Type I and Type II losses are rooted from the vector components of tangential and perpendicular electric field, respectively, whereas, the Type III losses corresponds the electric field complexity at the ends of the flat cell. The theoretical explanation provided in Hyde et al. states that the discontinuity in electric field around the sample surface due to the high dielectric constant of water results in the Q-value recovery and signal improvement[3]. The authors of Mett et al. also reported that arranging multiple flat cells in a stacked structure showed further improvement in signal[2].

As an extension to this study to a X-band cylindrical cavity, the authors of Sidabras et al. developed the aquastar cross-section of 3.5 *mm* O.D.[4]. These aquastar sample tubes were fabricated by the extrusion technique and the EPR signal intensity by a factor of 4 compared to a TPX 0.9 mm O.D. sample tube [4]. More complex cross-sections like the aquasun were also proposed in Sidabras et al., but the extrusion technique doesn’t support the fabrication of such complex designs[4].

Ultra or high-precision 3D printing has driven transformative advances in electronic, automotive, and healthcare industries as well as research. Stereolithography or SLA a form of additive manufacturing has gained wide traction in both research and industry due to its fabrication capabilities with micrometer-level precision. SLA is a layer by layer fabrication process, where a high definition ultraviolet light projector cures the photopolymer resin at fine resolution as small as 10 *µm*. Despite this inherent resolution, it is crucial to select the right process parameter and material. In this paper, we describe the approach for fabricating EPR sample tubes for X-band cavity and dielectric resonators, with emphasis the material selection, design optimization and printing parameters.

We designed, fabricated, and tested aquastar cross-sections with 1.5 *mm* O.D. for sapphire dielectric resonators. The 3D structures with the required cross-section were designed using ‘Autodesk Fusion 360’ and were fabricated using the ‘Boston Microfabrication (BMF) microArch S140’ with 10 *µm* resolution. The ‘Htl’ photopolymer resin used for 3D printing didn’t have any background signal, but the dielectric properties resulted in RF electric field losses. To lower the dielectric losses, we introduced the ‘latticing’ approach, where we removed excess plastic and introduced beams and pillars to hold the sample tube structurally sound. We utilized the high-resolution fabrication capabilities of the BMF 3D printers to print such complex designs at micrometer precision. The latticing approach resulted signal improvement in both simulations and experiments. In addition to the testing, we also present the optimizations on simulation to minimize computation time and also the 3D fabrication, to incorporate the error margin while printing. The signal calculation based on the theoretical approximation of lattice to a compound material(a mixture of vacuum and plastic at different compositions) based on the real and imaginary part of permittivity, significantly reduced the computation time within an error margin less than 0.5%.

The approach adopted in signal calculation for the aquastar cross-section for the dielectric resonator is extended to a complex cross-section aquasun in a cavity resonator, which permits more sample volume with lesser Q-value drop and improved signals. With the ultra-precision 3D printing, we developed aquasun sample tubes with high aspect ratio, which the extrusion technique doesn’t support. The novel aquasun geometries showed signal improvement both on simulations and experiments. Here in this paper, we present a simulation, fabrication, and optimization methodology in developing complex sample tube geometries.

## 2. Methods

### 2.1. Simulations

The finite element models were simulated on a Dell precision tower 7910 with dual Intel(R) Xeon(R) CPU E5-2670 @ 2.30 GHz processors and 512 GB DDR4 RAM with a Windows 11 operating system with 4 GB NVIDIA Quadro K4200 graphics card. The HFSS module on the Ansys Electronics Desktop version 2024 R1 was used to model the 9.5 *GHz* TE_01*δ*_ dielectric resonant cavity. Eigenmode solutions were used to model the magnetic field inside the cavities and determine the resonant frequency and the Q-value. The saturable (*Ss*) signal was calculated using the ‘Calculator’ option in the field overlays [5].

With the new sample holders, our aim is to improve the signal intensity of low concentration samples placed in an X-band dielectric resonator. The resonant cavity comprises a vacuum cylinder of 19.2 *mm* and a sapphire dielectric cylinder with an outer diameter of 8.2 *mm* and an inner diameter of 1.5 *mm* to place the sample tubes. Since the BMF (Boston Microfabrication) (10 *GHz ɛ* = 3.45 + *j* 0.084) [6] 3D printing material, ‘Htl’, the resin used to print the sample tubes was not in the material library, it was added as a custom material and the dielectric values, and the losses were defined manually. The Htl and PTFE (10 *GHz ɛ* = 2.1 + *j* 0.0021) were the sample tube materials used in our simulations. The anisotropic relative dielectric constant for the sapphire selected for the dielectric material is defined as 9.3, 9.3, and 11.6, along T(1,1), T(2,2), and T(3,3) axes respectively. The anisotropic dielectric loss tangent for the sapphire is defined as 3 *×* 10*^−^*^5^, 3 *×* 10*^−^*^5^, and 8.6 *×* 10*^−^*^5^, along T(1,1), T(2,2), and T(3,3) axes respectively. For every simulation, the sample space inside the sample tube was filled with water (10 *GHz ɛ* = 63 + *j* 29).

### 2.2. Slicing for computational efficiency

The 3D geometries designed using the Autodesk Fusion 360, while importing to ANSYS electronics desktop increased the computation time and observed design errors, due to the complex features in the design (mainly in latticed features). To reduce the computation time, the same geometries were again designed in the ANSYS HFSS for the simulation study. To further improve computational efficiency, the entire geometry was reduced to a thin slice of 1 *mm* and splitting the geometry along the vertical axis based on symmetry. For the sliced geometries, additional boundary conditions were added to the study to keep the resonant conditions same. The faces in the XY or the horizontal plane was assigned ‘PerfH’ boundary condition, which assumed the tangential component of the magnetic field to be zero. The cylindrical face of the cavity was assigned the ‘Finite Conductivity’, which considers the surface to have a specific electrical conductivity. In addition to this, same boundaries corresponding to the metal cavity was selected to calculate the associated power losses. To calculate the power losses from the sample, sample tube material, and dielectric resonator were defined in the model and the corresponding imaginary part of relative permittivity was defined in the calculator.

### 2.3. Sample Tube fabrication

The sample tube geometry was designed using the Autodesk Fusion 2024. The 3D geometry in .stl format was then exported for slicing to the VoxelDance additive software. The geometry was then sliced for 20 *µm* thickness at 10 *µm* resolution and each slice was converted to .png images which were then transferred to the BMF 3D printer.

The latticing was done using the ‘Volumetric Lattice’ extension in Autodesk Fusion. The geometry to be latticed was designed as a separate body in the same design, which was converted to the inbuilt basic lattice options (mainly gyroid and cross). The latticed geometry was converted to a mesh before exporting it for slicing. To ensure that the sample space inside the sample tube stayed leak proof and rigid, 50 *µm* wall was added in all lattice geometries. Since, the single exposure zone (projection size of the UV projector) have an area of 19.2 *mm* by 10.8 *mm*, approximately 40 sample tubes were printed in single batch with a fabrication time of 30 hours. After the completion of prints, the developed sample tubes were removed from the base layer and thoroughly cleaned using Isopropyl Alcohol (IPA) to remove any uncured resin. IPA was again pushed through the sample tube using a syringe to remove resin inside the sample space. In addition to this, the sample tubes were cured inside a UV chamber for 20 minutes, without heating to cure any remaining uncured resin.

### 2.4. Q-value measurement using VNA

The loaded Q-values of the sample tubes were determined by placing them in the dielectric resonator connected to the Keysight PNA-X network analyzer. The loaded Q-values were determined from the −3 dB bandwidth of the resonance. The quality factor extracted for an empty resonator was approximately 8000. To determine the loaded Q-values, the sample tubes were filled with water. The Q-values for all the sample tubes, along with the PTFE capillary were recorded using the VNA before the experimental measurements.

### 2.5. Signal Calculation

This section outlines the methodology used in determining the saturable and unsaturable signals from the CW-EPR spectra at X-band on an ELEXSYS 500 spectrometer (Bruker; Billerica, Massachusets) at room temperature with the following parameters: 345.8 *mT* center field, 10 *mT* scan width, variable incident microwave power, 14.68 *ms* conversion time, 60.13 second sweep time, 0.1 *mT* modulation amplitude, 100 *kHz* modulation frequency and 9.7 *GHz* microwave frequency. The printed sample tubes were filled with 100 *µM* TEMPOL solution in water and the EPR spectra were recorded at different microwave powers. The power saturation curves were then generated in Matlab by plotting the peak to peak signal intensity at the center field against the square root of microwave power. The saturable signal corresponds to the maximum value of peak to peak signal and the unsaturable signal was determined by finding the peak to peak signal when the square root of power was 0.6. The signal values were normalized to the 1.67 *µL* PTFE capillary.

## 3. Design Concepts

### 3.1. Aquastar Sample Tube Cross-Sections in a TE_01*δ*_ Dielectric Resonant Cavity

In Mett et al., the authors reported that placing flat cells perpendicular to the electric field nodal plane in a rectangular TE_102_ cavity reduces electric field losses[2]. Further improvements in the EPR saturable signal were observed by placing multiple flat cells in a similar orientation. As an extension to this methodology, in Sidabras et al., the aquastar cross-section with 3.5 *mm* O.D was developed for cavity resonators[4]. In Mett et al., authors extended the same aquastar cross-section with outer diameter 1.5 *mm*, which could be 3D printed and placed at the active region of the X-band sapphire dielectric resonator[7]. But, the 3D printing material used for developing the sample tubes have electric field losses due to the dielectric properties. In this article, we propose that the electric field losses could be reduced by removing the plastic and leaving only the 50 *µm* boundary wall(as in figure1A), minimal enough to hold the aqueous sample, but cannot be printed due to the structural instability of the design.

**Figure 1:**
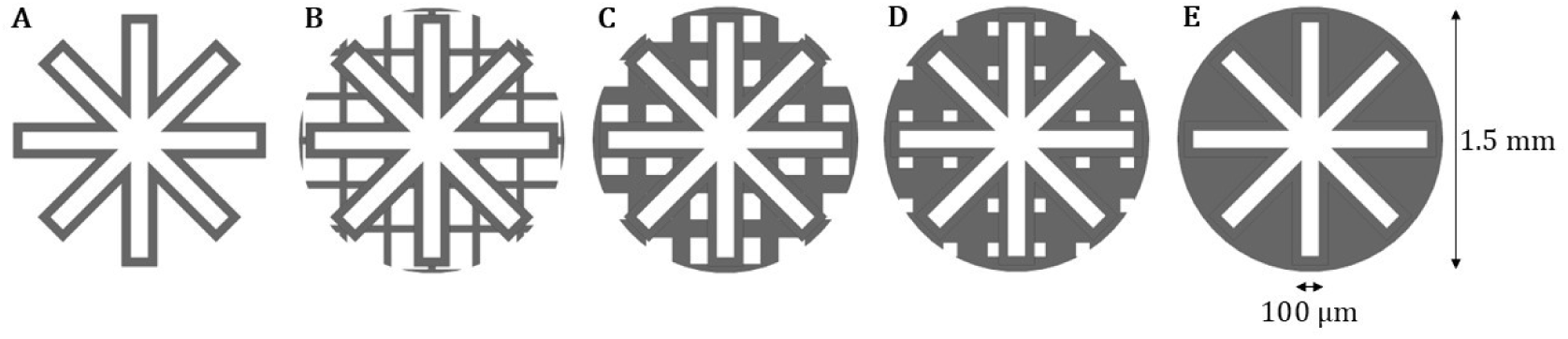
Cross-sectional views of BMF-printed Aquastar sample tubes with varying lattice solidities: (A) 0%, (B)20%, (C)40%, (D)60%, and (E)100%. The latticed cross-sections are designed to have a 50 *µm* boundary wall to hold the sample.

### 3.2. Latticing to minimize dielectric losses

In this paper, we have introduced the latticing approach in removing excess plastic, without compromising structural stability. Lattice structures mainly comprise of interconnecting beams and pillars, which significantly improves the stability by holding the thin structures sturdy, using minimum level of plastics. Since the BMF 3D printer could print at 10 *µm* resolution, we designed multiple lattice designs at different levels of plastic(figure1). For the experimental characterizations discussed in this paper, we printed gyroid and cross lattices at 20, 30 and 40 percentages of plastic. In addition to this, an aquastar sample tube with 100 percentage plastic was printed for comparison.

Since, latticed sample tubes had large number small faces, which increased the complexity of the simulations, cross lattices were designed in ANSYS. The lattices modeled for the simulations, were comprised of cylinders joined along different axes. To change the percentage of plastic, the radius of the cylinder was changed, corresponding to different solidity values. To model a different shape of lattice, the entire lattice structure was rotated 45^0^ by keeping one of the axes fixed. In addition to this, due to the complicated structure even after designing in ANSYS, slicing approach was adopted for the simulations, which significantly reduced the computation time. In addition to this, due to the complexity of designs, a custom meshing was applied for the simulations. The custom meshing defined a length-based high resolution meshing for the sample boundary wall and lattices as 0.285 *mm* and 0.3 *mm* respectively, whereas the rest of the volume had normal meshing.

### 3.3. Volume Approximation of Permittivity

Since the lattice structures comprised plastic and vacuum in different proportions, an additional simulation study was conducted based on the effective permittivity. In this study, the lattice volume was defined as a complete solid body, but the material was defined as custom material by changing permittivity based on the effective value of permittivity for plastic and vacuum mixture. The effective permittivity of the lattice was calculated based on three different mixing approaches: Simple volume Mixing, Bruggeman Mixing and Maxwell-Garnet Mixing [8][9][10].

The simple volume mixing approximates the effective permittivity *ε*_eff_:

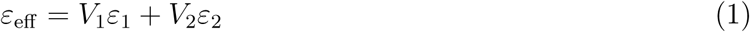

The Bruggeman mixing approximates the effective permittivity *ε*_eff_:

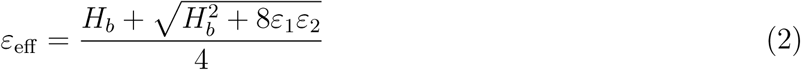

where, *H_b_* = (3*V*_2_ *−* 1)*ε*_2_ + (3*V*_1_ *−* 1)*ε*_1_.

The Maxwell-Garnet mixing approximates the effective permittivity *ε*_eff_:

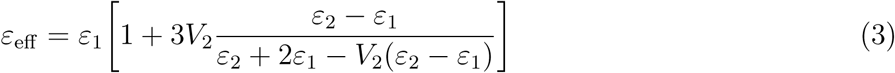

where, *ε*_1_, *ε*_2_, *V*_1_ and *V*_2_ = 1 *− V*_1_ correspond to the permittivity and fractional volumes of media 1 and 2 [11].

Based on the mixing rule, the effective permittivity was calculated using Matlab. The calculated values were entered to the permittivity and loss tangent sections of custom materials in the material properties for HFSS simulation. For this study, the fractional volumes were selected from 0 to 1 at 0.1 intervals, where 0 corresponds to 0% plastic and 1 corresponds to 100% plastic. The custom meshing defined in last section was adopted in this study for uniformity of results.

## 4. Results

### 4.1. Simulation Results

#### 4.1.1. Electromagnetic field distribution inside the Dielectric resonator

The electric and magnetic field distributions inside the resonator, calculated using the eigenmode studies are given in figure 2. The field distribution inside the dielectric resonator showed concentrated homogeneous magnetic field at the center of the cavity, and the electric field resides in sapphire due to its dielectric properties. The magnetic field shown in figure 2 was 90 degrees out of phase compared to the electric field. The model of the empty cavity before placing the sample and the sample tube had an approximate Q-value of 7000.

**Figure 2:**
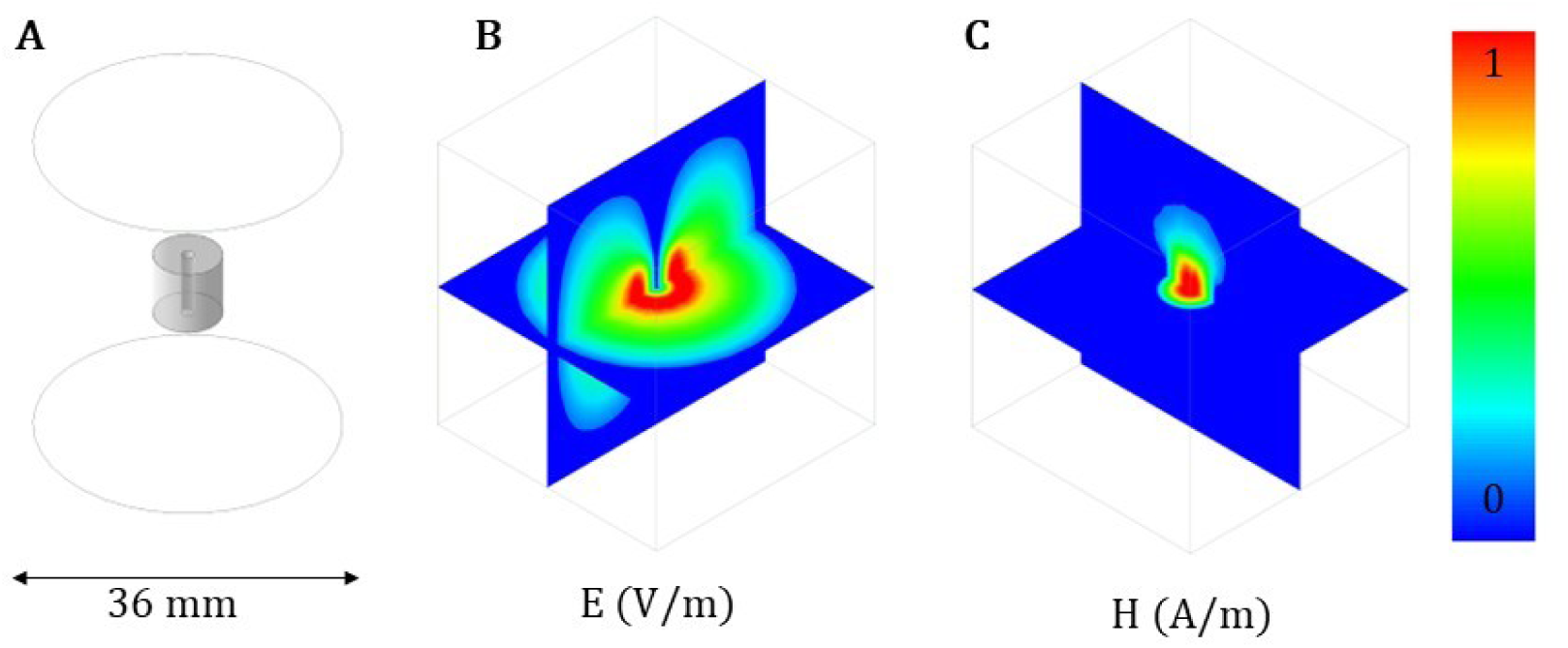
(A)Dielectric resonant cavity selected for the simulations, (B)Electric, and (C)Magnetic field distribution inside the resonator.

#### 4.1.2. Signals calculated from Simulations

For each sample tube configuration, the saturable EPR signal strength or saturable signal (*Ss*) was calculated with the HFSS fields calculator using the methods and equations mentioned in Mett et al.[12]. The equation for saturable signals (*Ss*) is given as:

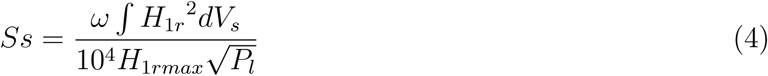

where *ω* corresponds to the resonant frequency, *H*_1*r*_ is the rotating RF magnetic field component in the sample and max indicates the maximum value of the sample volume, *V_s_* represents the sample volume and *P_l_* represents the total power losses, which includes the metallic walls, dielectric, sample, and sample tube.

The percentage of plastic was varied by changing the radius of lattice using the parametric study, and the signal values were calculated for each radius value. The normalized signals determined from the simulation are plotted against the solidity level and are given in figure 4. The plot showed a clear inward relation of signal to level of plastic, suggesting the increased dielectric losses with high solidity.

**Figure 3:**
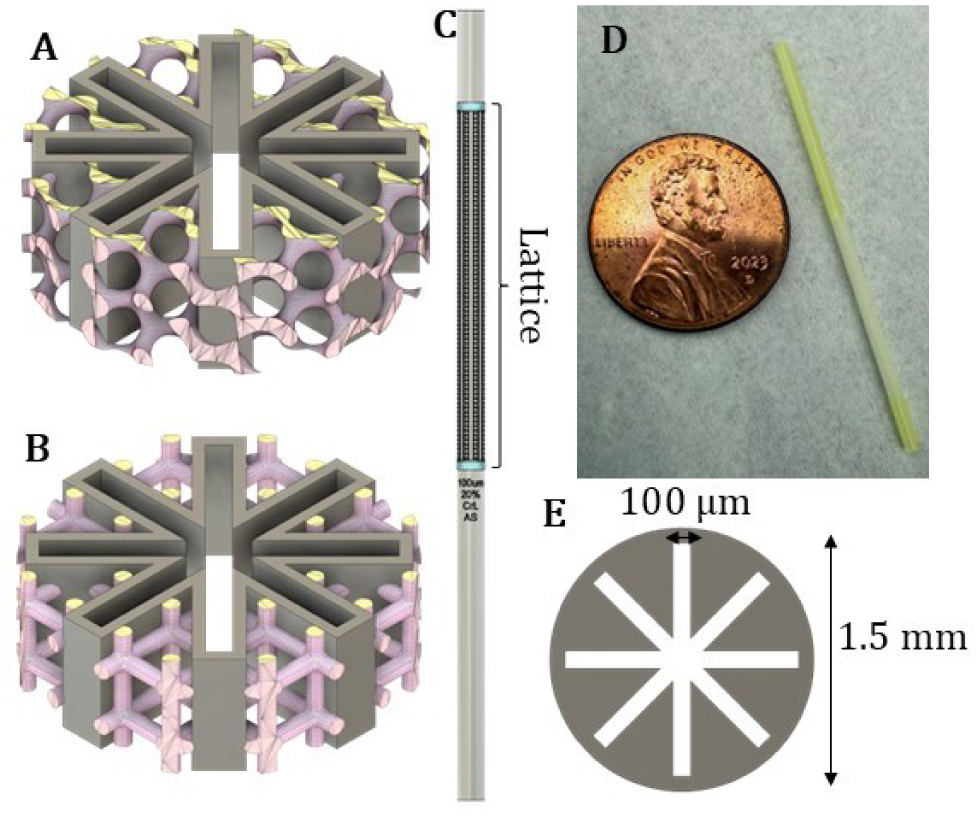
Sample tube designs. (A)20% gyroid lattice and (B)20% cross lattice. (C)3D model of a 20% latticed sample tube; (D) 3D-printed sample tube and (E)Aquastar cross-section. [7, 4]

**Figure 4:**
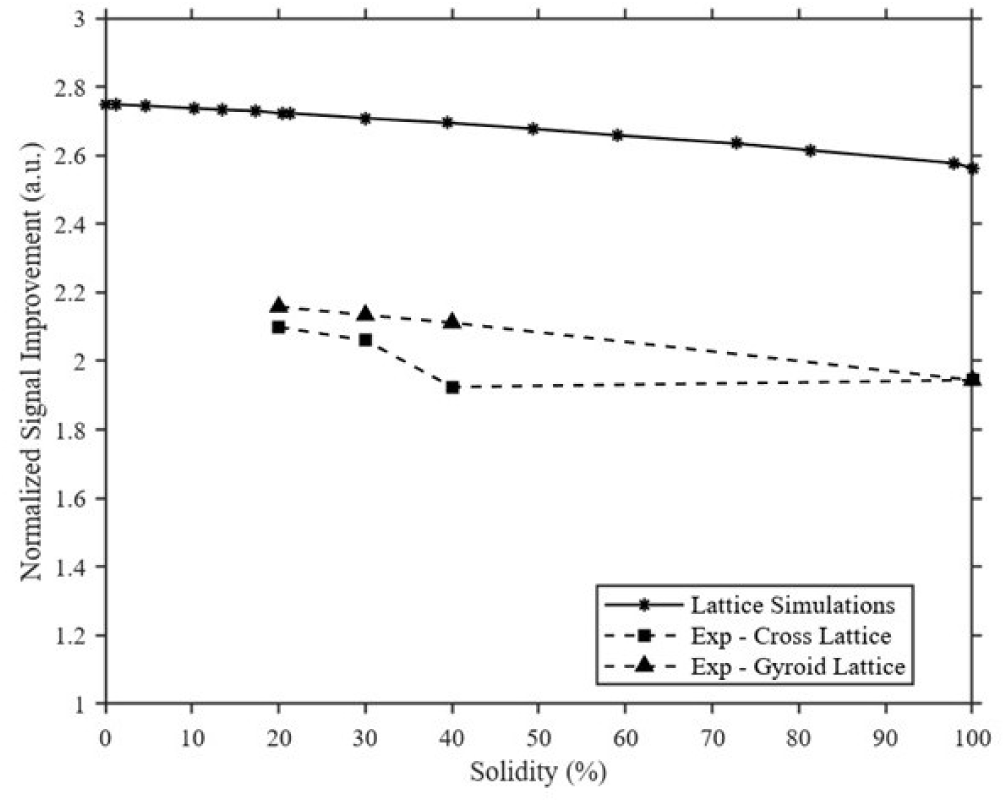
Simulated and experimental signals plotted against solidity. The signals are normalized to a PTFE capillary sample tube.

**Figure 5:**
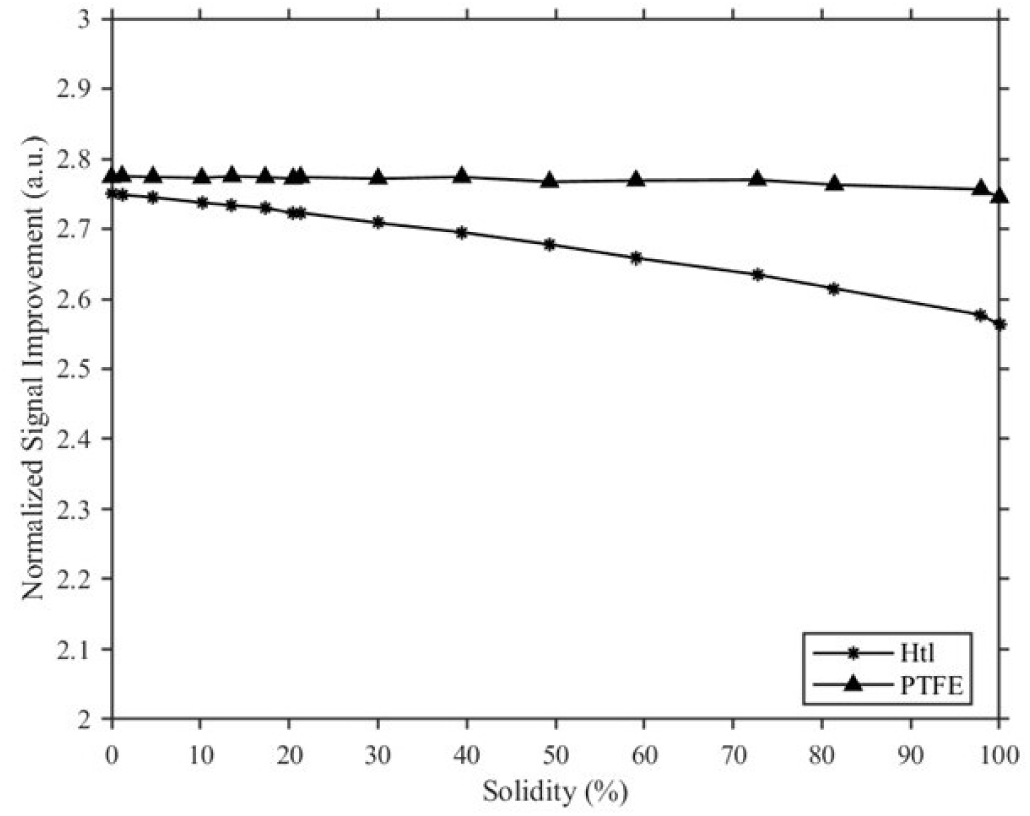
Normalized signal plotted against solidity for Htl and PTFE materials.

**Figure 6:**
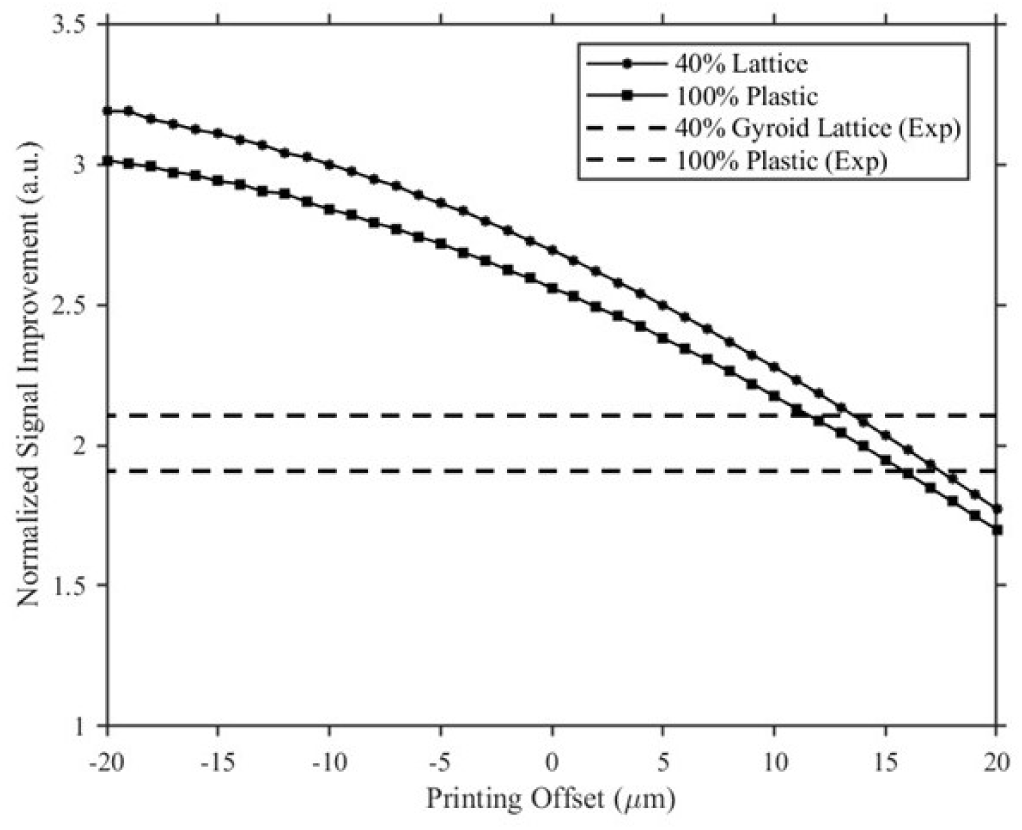
Printing optimization results. Normalized signal plotted against the printing offset. The dash lines represent the experimental results.

As mentioned in the methods section, since we printed cross and gyroid lattices for our experiments as in figure 3 A and B, but the same designs couldn’t be imported due to complex features in the design. To investigate the significance of lattice shape in signal, we followed an indirect approach in changing the lattice shape, by tilting the entire lattice. The normal and tilted lattice slices and the signals plotted against the solidity is given in supplementary figure 14. The signals didn’t show any dependence on the lattice shape but still follow the inward relation with volume of plastic.

In addition to the above studies, we used the same latticing approach to determine the signals if the sample tube material was replaced with teflon instead of Htl. The signal to solidity plot determined from this study is given in figure 5. The plot showed further improvement in signal for teflon due to their excellent dielectric properties while following the same inward relation with the volume of plastic.

### 4.2. Q-value Versus Percentage of Plastic

The resonant frequencies and Q-value measured using the network analyzer are given in the table 1. Since UV curing is an important procedure during the sample tube fabrication, the Q-values were measured before and after UV exposure. The increase in quality factor for both the gyroid and cross lattices suggest the decrease in dielectric losses with the reduction of plastic. The Q-value didn’t show a drastic change after the UV curing shows that there wasn’t any uncured resin left after cleaning.

**Table 1:** Resonant frequency and Q-value measured before and after curing.

| Sl. No. | Sample Tube Design | Before curing |  | 20 mins curing |  |
|---|---|---|---|---|---|
|  |  | Frequency (GHz) | Q-value | Frequency (GHz) | Q-value |
| 1 | Empty Cavity | 9.709 | 7550 | – | – |
| 2 | PTFE 1.5 mm OD | 9.706 | 3101 | – | – |
| 3 | 100% Plastic | 9.705 | 2995 | 9.705 | 2951 |
| 4 | 40% Gyroid Lattice | 9.706 | 3592 | 9.706 | 4003 |
| 5 | 30% Gyroid Lattice | 9.707 | 4006 | 9.706 | 4090 |
| 6 | 20% Gyroid Lattice | 9.707 | 4164 | 9.707 | 4172 |
| 7 | 40% Cross Lattice | 9.705 | 3203 | 9.706 | 3728 |
| 8 | 30% Cross Lattice | 9.706 | 3575 | 9.706 | 3820 |
| 9 | 20% Cross Lattice | 9.707 | 3909 | 9.707 | 4176 |

### 4.3. Signal Versus Percentage of Plastic

This section comprises the saturable signal (Ss) calculated from the experiments. The CW-EPR spectra for each sample tube recorded at a microwave power of 0.7962 mW is given in supplementary figures 12 and 13. The spectra showed a 2-fold improvement for the 3D-printed aquastar compared to the PTFE capillary. In addition to this, the latticed sample tubes showed further improvements for both lattice types. The power saturation curves were fitted based on the first derivative peak to peak amplitude versus power fit equation given in Altenbach et al.[13]. The *P*_1*/*2_ were calculated from the power saturation fit and were normalized to the PTFE capillary and are given in figure 4 and table 2. The calculated Ss is given in table 2 and are plotted against solidity along with the simulations in figure 7. To determine the effect of lattice on the resonator efficiency or *λ*_Sim_ were calculated based from the simulations and are given in table 2. The experimental results showed approximately 20% deviation compared to simulations, which roots mainly from the printing errors and this is discussed in detail in the discussion section.

**Figure 7:**
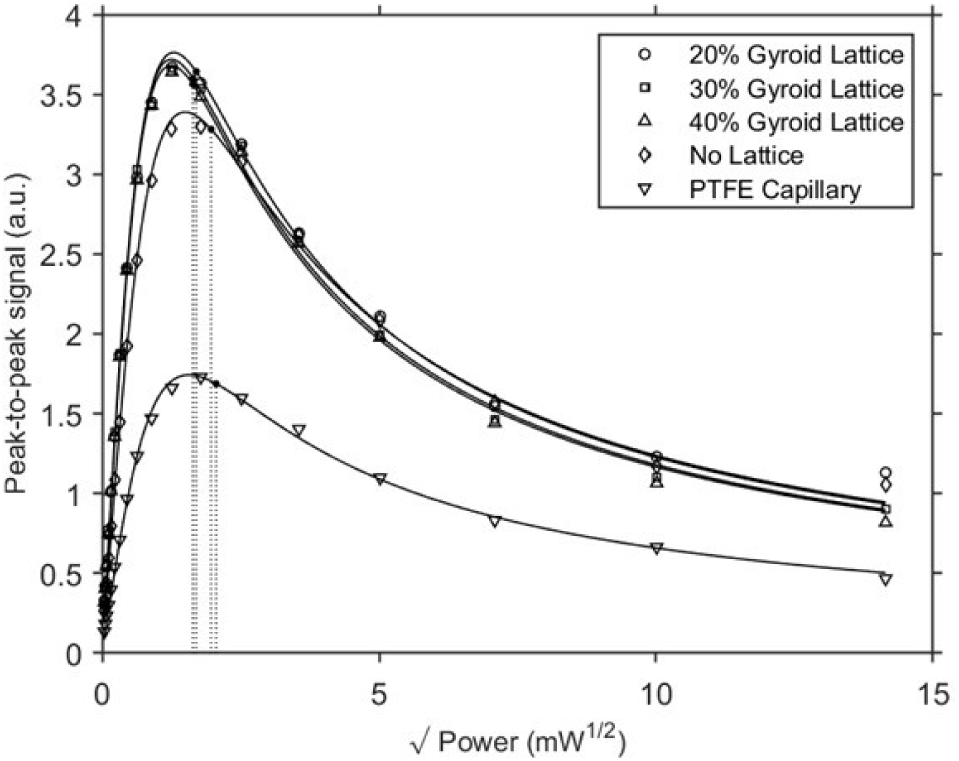
Experimental results of power saturation measurements for gyroid-type lattice sample tubes. Peak to peak signal plotted against square root of power and fitted to power saturation fit. Each measurement was repeated (n = 2) for reproducibility.

**Table 2:**
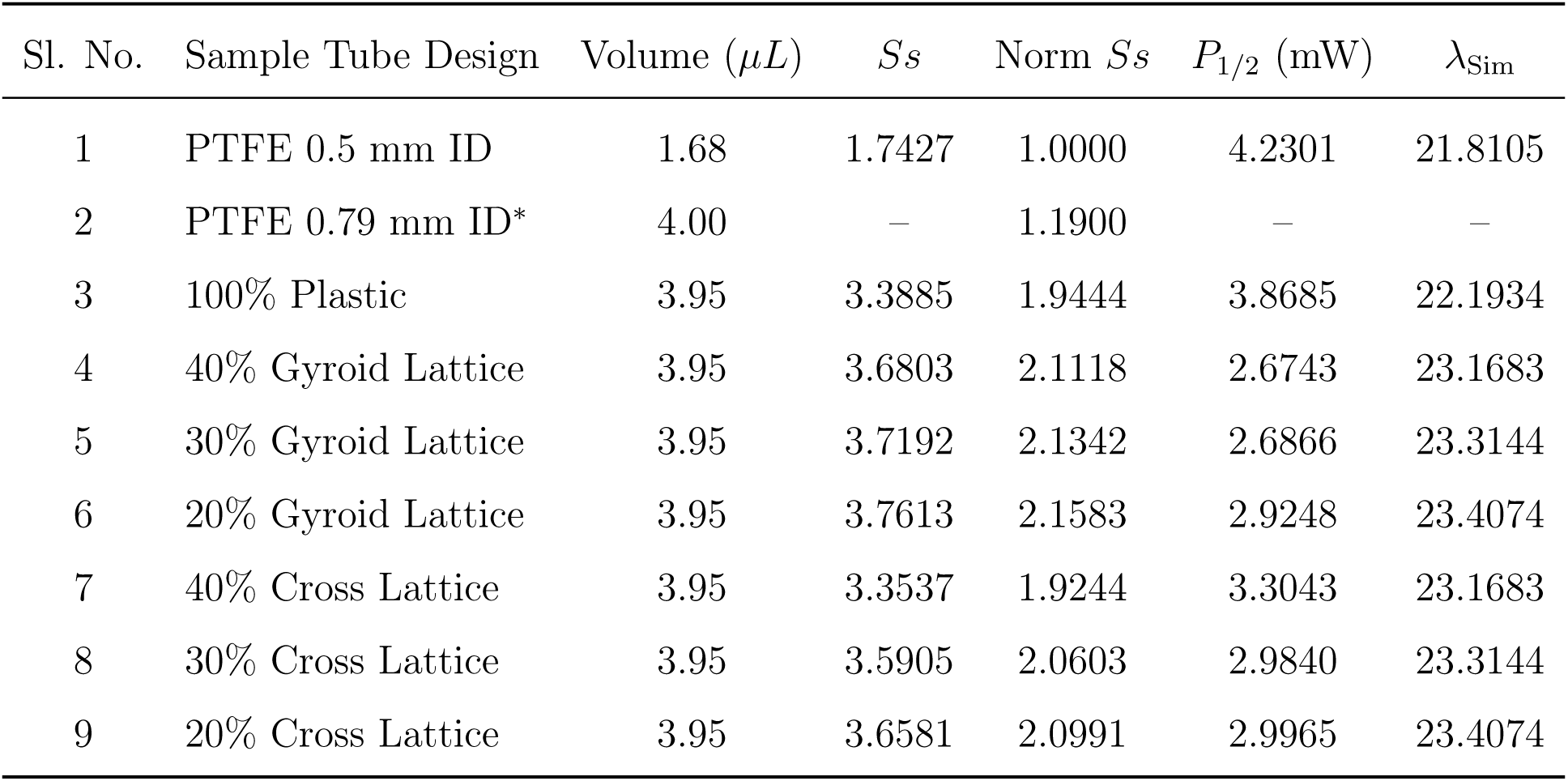
Signal normalization for different sample tube designs.

| Sl. No. | Sample Tube Design | Volume (*µL*) | *Ss* | Norm *Ss* | *P*<sub>1/2</sub> (mW) | λ<sub>Sim</sub> |
|---|---|---|---|---|---|---|
| 1 | PTFE 0.5 mm ID | 1.68 | 1.7427 | 1.0000 | 4.2301 | 21.8105 |
| 2 | PTFE 0.79 mm ID* | 4.00 | – | 1.1900 | – | – |
| 3 | 100% Plastic | 3.95 | 3.3885 | 1.9444 | 3.8685 | 22.1934 |
| 4 | 40% Gyroid Lattice | 3.95 | 3.6803 | 2.1118 | 2.6743 | 23.1683 |
| 5 | 30% Gyroid Lattice | 3.95 | 3.7192 | 2.1342 | 2.6866 | 23.3144 |
| 6 | 20% Gyroid Lattice | 3.95 | 3.7613 | 2.1583 | 2.9248 | 23.4074 |
| 7 | 40% Cross Lattice | 3.95 | 3.3537 | 1.9244 | 3.3043 | 23.1683 |
| 8 | 30% Cross Lattice | 3.95 | 3.5905 | 2.0603 | 2.9840 | 23.3144 |
| 9 | 20% Cross Lattice | 3.95 | 3.6581 | 2.0991 | 2.9965 | 23.4074 |

**Table 3:**
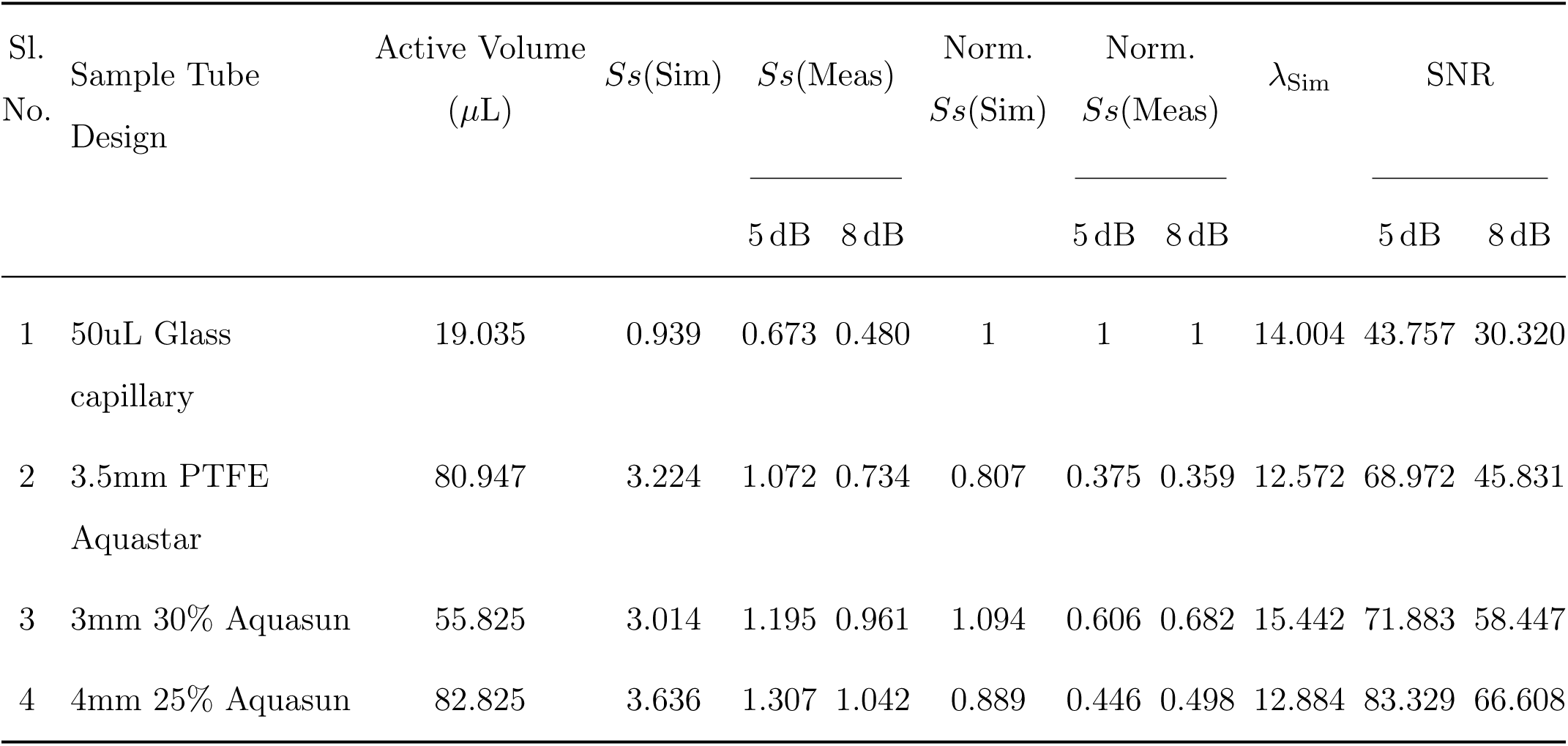
Signal normalization for Aquasun sample tube designs. The aquasun designs were placed in a NMR capillary of appropriate size and the same was considered during the simulations too. The power saturation experiments were conducted using aqueous 30uM T4L118R1 solution. The normalized signals represent the simulated and measured <u>signals normalized based on volume.</u>

| Sl.<br>No. | Sample Tube Design | Active Volume<br>(µL) | *Ss*(Sim) | *Ss*(Meas) |  | Norm.<br>*Ss*(Sim) | Norm.<br>*Ss*(Meas) |  | λ<sub>Sim</sub> | SNR |  |
|---|---|---|---|---|---|---|---|---|---|---|---|
|  |  |  |  | 5 dB | 8 dB |  | 5 dB | 8 dB |  | 5 dB | 8 dB |
| 1 | 50uL Glass capillary | 19.035 | 0.939 | 0.673 | 0.480 | 1 | 1 | 1 | 14.004 | 43.757 | 30.320 |
| 2 | 3.5mm PTFE Aquastar | 80.947 | 3.224 | 1.072 | 0.734 | 0.807 | 0.375 | 0.359 | 12.572 | 68.972 | 45.831 |
| 3 | 3mm 30% Aquasun | 55.825 | 3.014 | 1.195 | 0.961 | 1.094 | 0.606 | 0.682 | 15.442 | 71.883 | 58.447 |
| 4 | 4mm 25% Aquasun | 82.825 | 3.636 | 1.307 | 1.042 | 0.889 | 0.446 | 0.498 | 12.884 | 83.329 | 66.608 |

### 4.4. Volume Approximation of Permittivity

The results obtained from the volume approximation of permittivity are presented in this section. The normalized signal calculated for each lattice to volume ratio is plotted against the percentage of plastic along with the simulated results for lattices and are given in 9. The signals calculated from the permittivity approximations were in excellent agreement with the simulations with an approximate error margin less than 1%. The Maxwell-Garnet mixing rule approximation yielded the minimum deviation with a 0.3% error margin.

**Figure 8:**
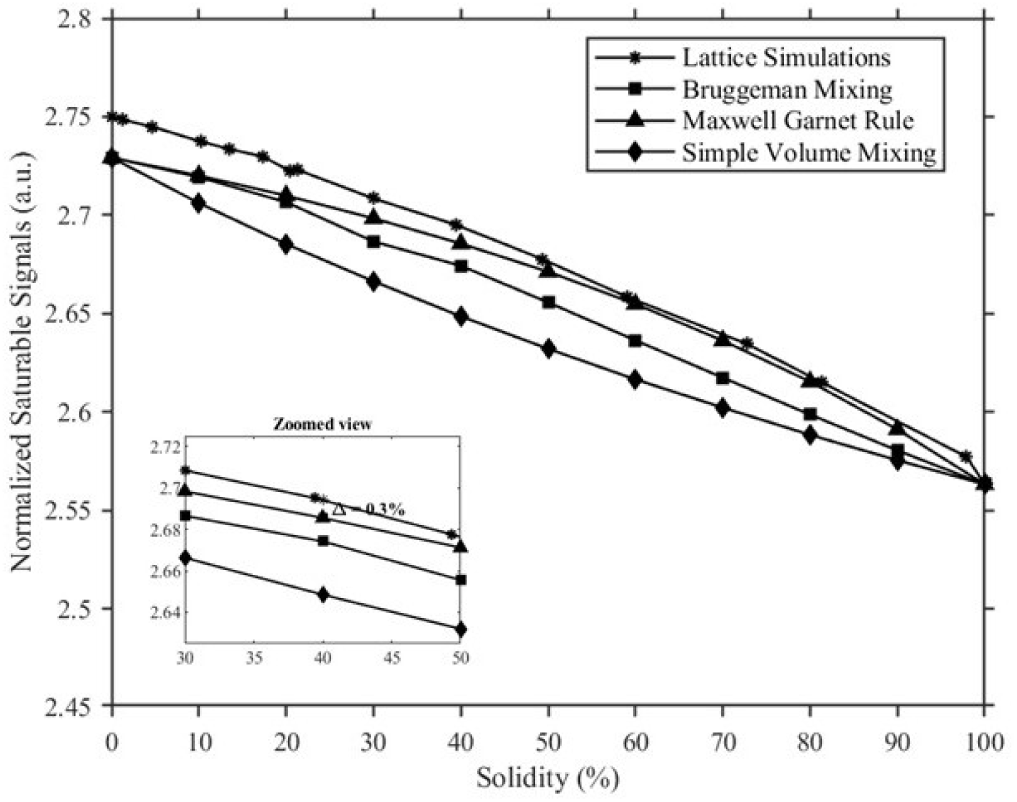
Normalized signal determined from the dielectric approximation study plotted against solidity.

## 5. Discussion

### 5.1. Printing optimization for better signals

The signal to percentage of plastic plot followed an inward relation for all the experimental and simulation studies. It should also be noted that the experimental results showed approximately 20% loss in signals, which was mainly due to the error margin in the resolution of the BMF 3D printer. We conducted an additional study to investigate this offset in printing. The offset in printing was defined as a parameter and was swept from −20 *µm* to 20 *µm* at 1 *µm* increments, while keeping the solidity constant at 40% and 100%. The calculated signals plotted against the printing offset is given in figure 6. By comparing the experimental results, a printing error margin of 13 *µm* to 18 *µm* was determined from this study. These values should be accommodated while designing the sample tubes to achieve better accuracy in experiments to simulations, as 3D printers inherently have an error margin.

### 5.2. Volume Approximation of Permittivity

The slicing approach defined in the methods section significantly reduces computation time and complexity. As a further improvement in computational cost, the volume approximation of permittivity approach is described in the methods section. The volume approximation approach significantly reduced the total number of elements in the study from approximately 800,000 to around 7000, with minimum deviation. This approach is mainly helpful in computing the signals in latticed sample tubes. From the dielectric study discussed in the methods and results discussion section, it was observed that the signal values corresponding to the ‘Maxwell-Garnet mixing’ rule were closer to the simulations compared to the ‘Bruggeman mixing’ and ‘simple volume mixing’ rules.

### 5.3. Latticing approach for complex cross-sections

The latticing approach in reducing the amount of plastics is crucial in complex cross sections like the ‘Aquasun’ geometry designed for cavity resonators (figure 11D). The aquasun cross-section is an extension of the aquastar geometry, with increased sample volume with minimum dielectric losses. The seed idea for this geometry came form the Sidabras et al. 2017, where the authors proposed the simulation study of a 10 mm O.D. sample tube with 4 main fins and 12 sub fins arrangement. Till date, the PTFE extrusion technique doesn’t support the fabrication of such geometries due to the wide aspect ratio difference, But with the ultra-precision 3D printing, these designs are fabricated using the BMF-Htl material. But the crucial thing to be considered here is the dielectric properties of the photopolymer material. The latticing approach reduces the dielectric losses to an extend, but having such high sample volumes inside the cavity significantly lower the Q-values and tuning was extremely difficult. Upon multiple trials, we finalized 3 mm, 4 mm, and 5 mm O.D. aquasun geometries with 0.5 I.D, inner capillary, and 8 separate fins surrounding the inner capillary. These novel sample tubes presented higher signals compared to the capillary and aquastar geometries. The detailed study of these cross-sections including both simulation and experimental verification is discussed here in this section.

**Figure 9:**
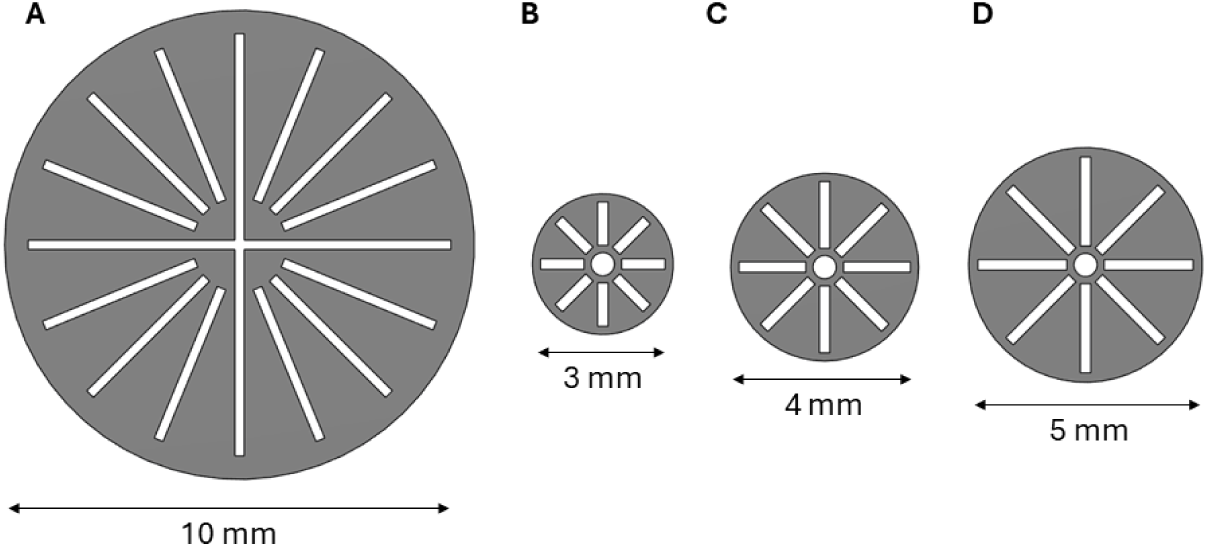
Cross-sectional images: (A) 10 mm Aquasun propsed in Sidabras et al. 2017, (B) 3 mm, (C) 4 mm, and (D) 5 mm Aquasun cross-sections selected for 3D printing. Designs given in panel (B), (C), and (D) have 225um fins, and a central capillary of 0.5 mm I.D.

In Sidabras et al. 2017, the authors presented the 10 mm O.D aquasun cross-sections were simulated, but the extrusion fabrication using teflon didn’t support it’s development due to the higher outer diameter to fin width ratio comparing to aquastar cross-section. By introducing ultra-precision 3D fabrication, we are now able to develop cross-sections at the proposed resolution, even with complex designs. The crucial thing to consider while 3D printing the aquasun designs with higher amount of plastic is to consider the dielectric properties of the 3D printing photopolymer. Since, these designs have a higher amount of plastic compared to the aquastar designs, the latticing approach is effective in reducing plastic, thereby reducing dielectric losses. The aquasun designs were simulated through the slicing approach discussed in the methods section. We sliced the geometry along the perpendicular planes, making it quarter of the slice, and applying the ‘PerfE’ boundary conditions and instead of designing the latticed geometry, we followed the dielectric approximation method, which further reduced the computation time. The signals calculated for the aquasun were normalized to 0.9 mm OD glass capillary.

**Figure 10:**
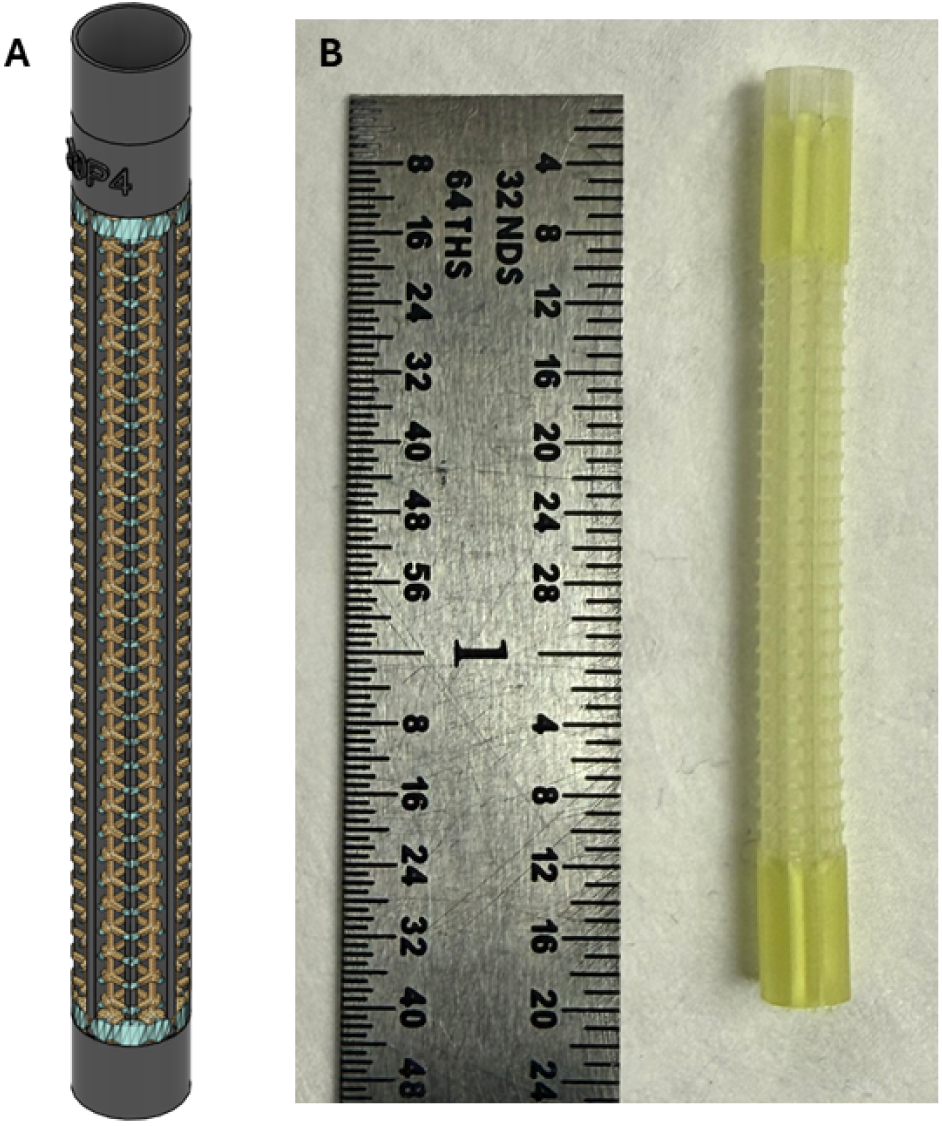
(A) Perspective view of 4 mm aquasun sample tube designed for cavity resonator. (B) Fabricated Aquasun sample tube.

The aquasun sample tubes were designed using the Autodesk Fusion 360. The sample tubes were designed with open cross-sections at both ends, with bucket on top to load the sample. A 25 mm latticing region was kept in the middle to have maximum overlap of the TE011 magnetic field. Once printed, the sample tubes were thoroughly cleaned multiple times using an IPA stream to remove any uncured resins. The sample tubes were then cured in a UV chamber for 90 minutes. The Q-value and the frequencies were noted in a VNA every 30 minutes and an inward relation between the amount of plastic and the Q-value were observed. As these designs also have higher volume of plastic compared to the 1.5 mm aquastar designs, a significant improvement in Q-value was observed before and after curing. Since the Q-value didn’t profoundly increase from 60 minutes to 90 minutes curing, we finalized 90 minutes of UV exposure time is chosen for our further fabrications.

Due to the material properties and the rough surfaces created due to additive manufacturing, the simple capillary action didn’t fill the sample tubes. To efficiently fill the samples, we sealed the bottom end of the sample tube with citriseal and loaded the sample in the top bucket. These top loaded sample tubes were then placed in vacuum chamber, which pulls out the air bubbles and the aqueous sample replace them, once the vacuum chamber is back to atmospheric pressure. This sample loading technique was repeated a few times to make sure the latticed region is fully filled. The Q-values and the frequencies for the loaded sample tubes were determined before the power saturation experiments.

During the power saturation experiments, we observed a TEMPOL quenching, possibly the photopolymer material property, which diminished the signal intensity over time (which was experimentally verified and is given in the supplementary section along with the power saturation results), we used a spin labeled aqueous protein sample (R1 labeled T4 Lyzosome, T4L 118R1). The spectra recorded for the T4L118R1 is given in figure 11. The deviation of the simulated signals from the power saturation results lead us to verify the dielectric properties of the photopolymer material. To experimentally determine the relative permittivity values, we followed ASTM 2520 protocol, where a 3D printed cylindrical block of Htl material was inserted to determine the shift in resonant frequency and Q-value shifts and thereby determine the real and imaginary components of relative permittivity. The experimentally determined values were slightly different from the published datasheet values, and the determined values are given in supplementary table.

**Figure 11:**
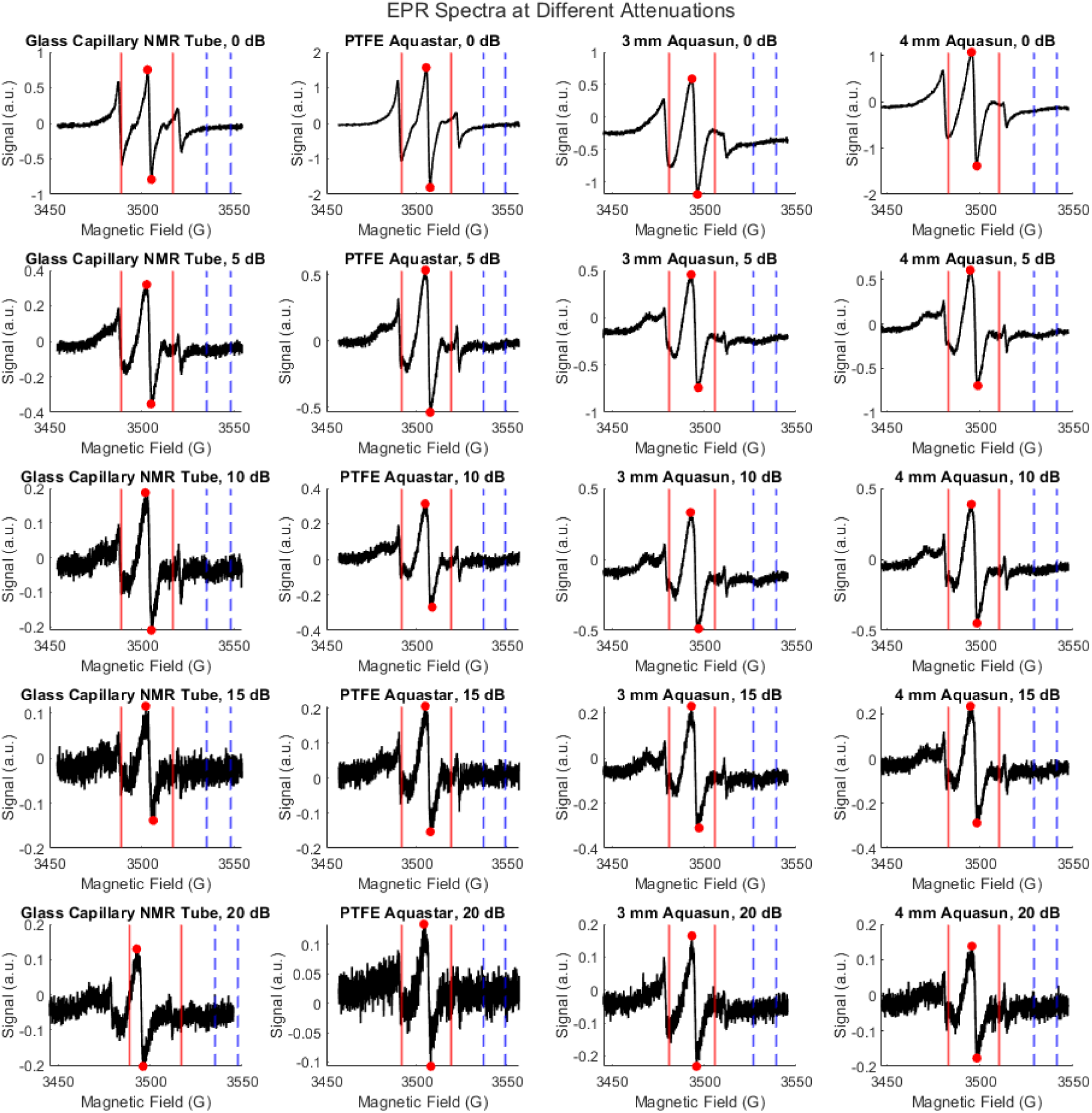
Representative CWEPR spectra of aqueous T4L118R1 solution loaded in different sample tubes and at different microwave attenuation. The interval inside the blue dashed line represent the field interval used for SNR calculation. and the red lines represent the field interval used for peak to peak intensity calculation.

To verify the experimental results, the sample tubes used for testing were simulated with the experimentally determined dielectric values. We followed the dielectric approximation methodology based on ‘Maxwwell-Garnet mixing’ discussed earlier, in simulating the lattices with minimum computation time. The 3D printed aquasun sample tubes have signal improvement greater than or equal to the PTFE aquastar sample tubes both in simulations and experiments. The signal intensities normalized to sample volume showed significant improvement compared to the PTFE aquastar. Even though, simulated and the experimental results show slight deviation, the normalized signals shows significant improvement compared to the PTFE aquastar. It should also be noted that the SNR (signal to noise ratio) is improved for the aquasun sample tubes the compared to both glass capillary and the PTFE aquastar sample tubes. Here, with the discussed findings and methodology, sample tubes can be developed based on the material properties.

## 6. Conclusion

With this project, we demonstrated the application of ultra-precision 3D printing in fabricating complex-sections and optimizing the designs to maximize the signals. The aquastar sample tubes were printed using the BMF MicroArch S140 at 10 *µm* resolution and had an active volume of approximately 3-4 *µL*. The printing and post-processing time for a sample tube was approximately 2 hours (calculated based on the batch fabrication process). It was observed from both the simulations and experiments that removal of plastic reduces the dielectric losses, and thereby improving the signals, without compromising structural integrity. The approach in designing the latticed sample tubes is discussed in-depth in the methods section. The signals calculated from the power saturation experiments showed approximately 20% deviation from the simulations, which we concluded mainly originated from the printing error. This error was determined from the simulations and was observed around 13 *µm* to 18 *µm*. From the plots for signals versus solidity for PTFE material, showed further improvement due to the dielectric properties. The slicing approach based on symmetry discussed in this study reduced computation time mainly for latticed structures. In addition to this, we determined that the lattice shape doesn’t have any effect in the signal. As an extension to this, we conducted the volume approximation of permittivity study, which further reduced the computation time and the calculated signals were close to the findings from the simulations.

The design and simulation methodology and the findings are extended to develop the complex aquasun cross-sections for cavity resonators. With the slicing approach and

In summary, we have showed an efficient method to fabricate sample tubes using 3D printing, while accounting the dielectric losses in the design. The demonstrated method increases the spin sensitivity particularly for low concentration samples. In addition to this, by discussing the approach and findings from the aquasun cross-section, we open the questions to further development of various aqueous sample configurations.

## 7. Acknowledgment

The authors thank Boston Micro Fabrication (Marynard, MA) for the 3D printer support. This work was supported by the National Institutes of Health, National Institute of General Medical Sciences (NIH, NIGMS) grant GMA149568 (to JWS).

## 8. Supplementary Section

### 8.1. TEMPOL quenching in aquasun cross-section

It was observed during the power-saturation experiments using aquasun geometry, that the TEMPOL peak to peak signal diminishes over time. It is to be noted that the sample volume didn’t reduce, ruling out the sample leaking, and suggesting a quenching of TEMPOL by the 3D printing photopolymer. To verify this, we conducted a time-delayed overnight spectrum, where we took CWEPR spectrum in 5 minutes interval over 8 hours. The TEMPOL quenching was not so evident in the smaller aquastar geometries, as they had lower surface area, compared to the bigger aquasun geometries.

**Figure 12:**
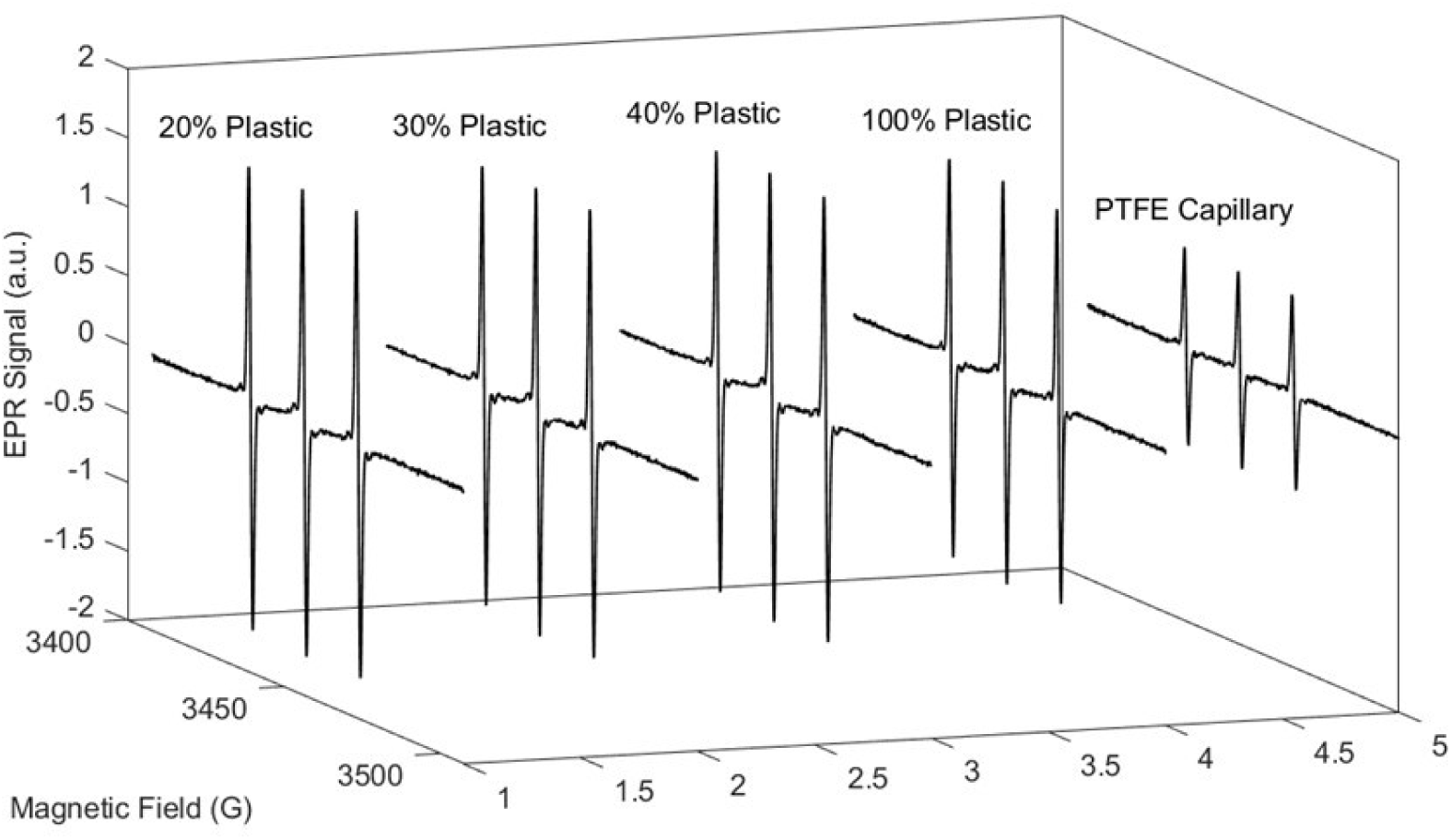
CW-EPR spectra for 100 *µM* aqueous TEMPOL solution recorded at 0.7962 *mW* for 20%, 30%, 40%, 100% plastics and PTFE capillary plotted for Gyroid type lattice.

**Figure 13:**
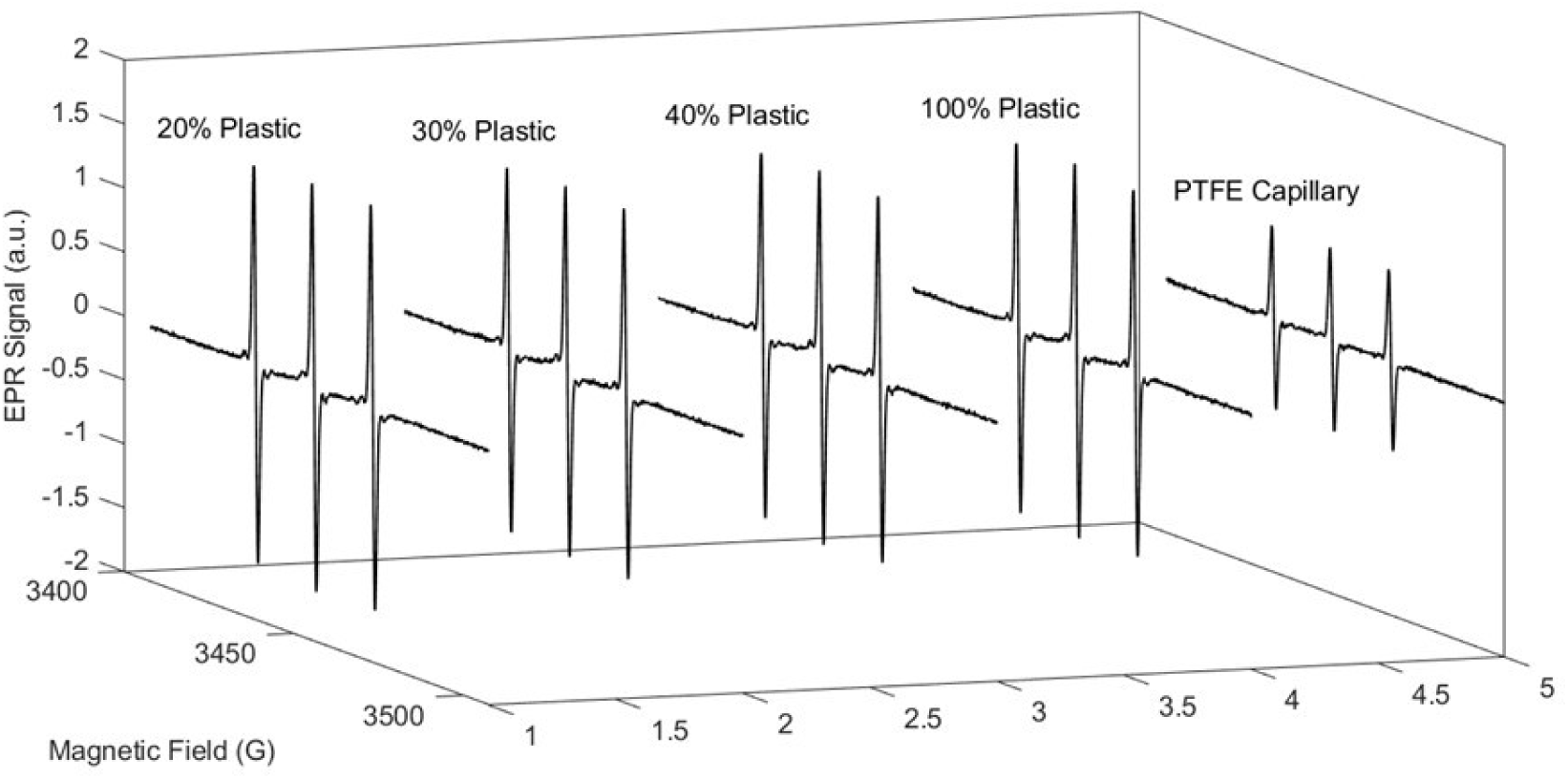
CW-EPR spectra for 100 *µM* aqueous TEMPOL solution recorded at 0.7962 *mW* for 20%, 30%, 40%, 100% plastics and PTFE capillary plotted for cross type lattice.

**Table 4:** Signal normalization for Aquasun sample tube designs. The 3mm aquasun sample tube was placed in a NMR capillary of appropriate size and the same was considered during the simulations too. The power saturation experiments were conducted using aqueous 180uM TEMPOL solution. The normalized signals represent the simulated <u>and measured signals normalized based on volume.</u>

| Sl. No. | Sample Tube Design | Active Volume (*µ*L) | *Ss*(Sim) | *Ss*(Meas) |  | Norm. *Ss*(Sim) | Norm. *Ss*(Meas) |  | λ<sub>Sim</sub> | SNR |  |
|---|---|---|---|---|---|---|---|---|---|---|---|
|  |  |  |  | 5 dB | 8 dB |  | 5 dB | 8 dB |  | 5 dB | 8 dB |
| 1 | 50uL Glass capillary | 19.035 | 0.939 | 0.673 | 0.480 | 1 | 1 | 1 | 14.004 | 43.757 | 30.320 |
| 2 | 3.5mm PTFE Aquastar | 80.947 | 3.224 | 1.072 | 0.734 | 0.807 | 0.566 | 0.528 | 12.572 | 68.972 | 45.831 |
| 3 | 3mm 30% Aquasun | 55.825 | 3.014 | 1.195 | 0.961 | 1.094 | 0.660 | 0.759 | 15.442 | 71.883 | 58.447 |
| 4 | 5mm 25% Aquasun | 108.475 | 3.795 | 1.307 | 1.042 | 0.709 | 0.364 | 0.369 | 12.884 | 83.329 | 66.608 |

**Figure 14:**
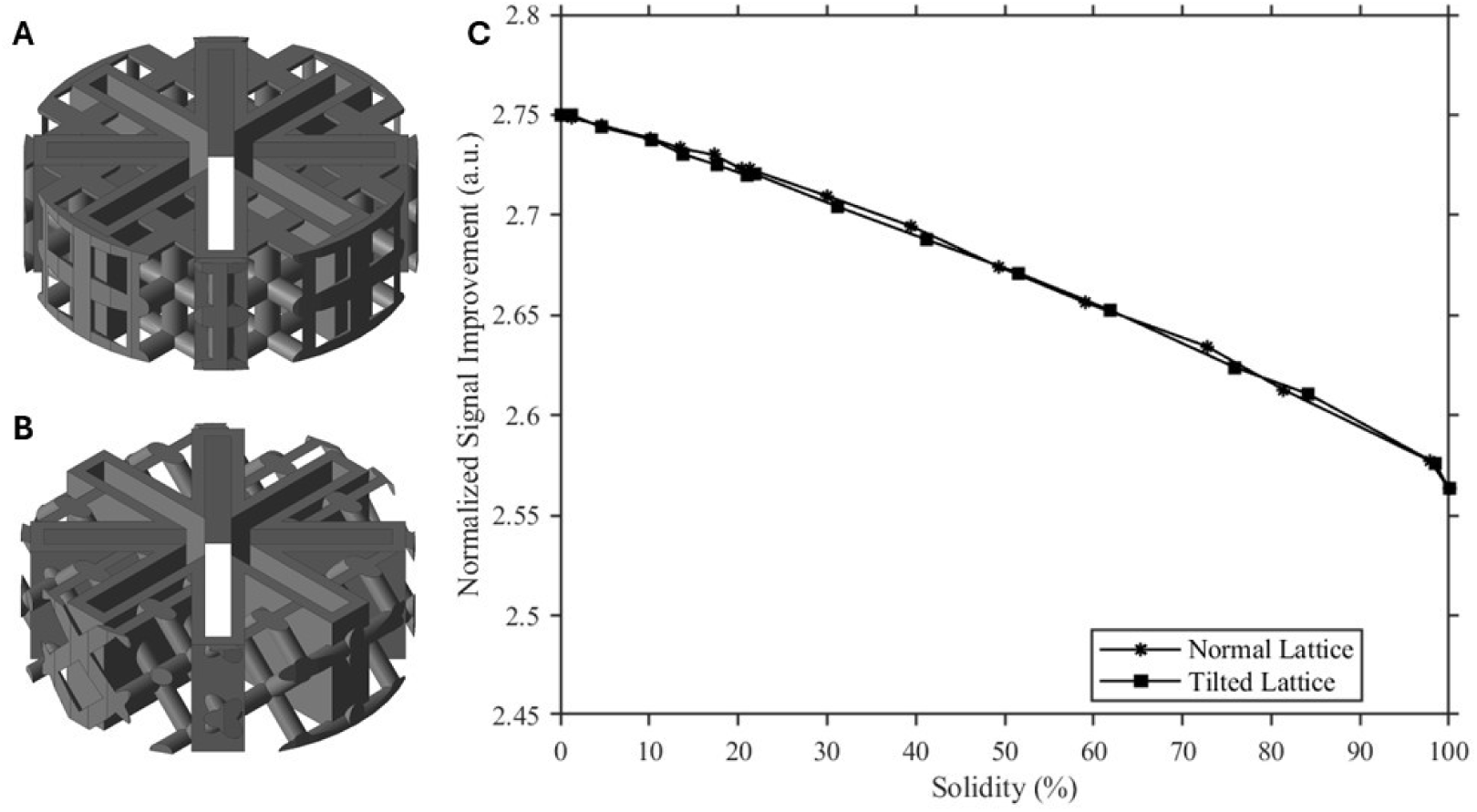
Lattice shape dependence on signal: (A)Normal lattice, (B)Tilted lattice, and (C)Normalized signal plotted against solidity for normal and tilted lattices.

**Figure 15:**
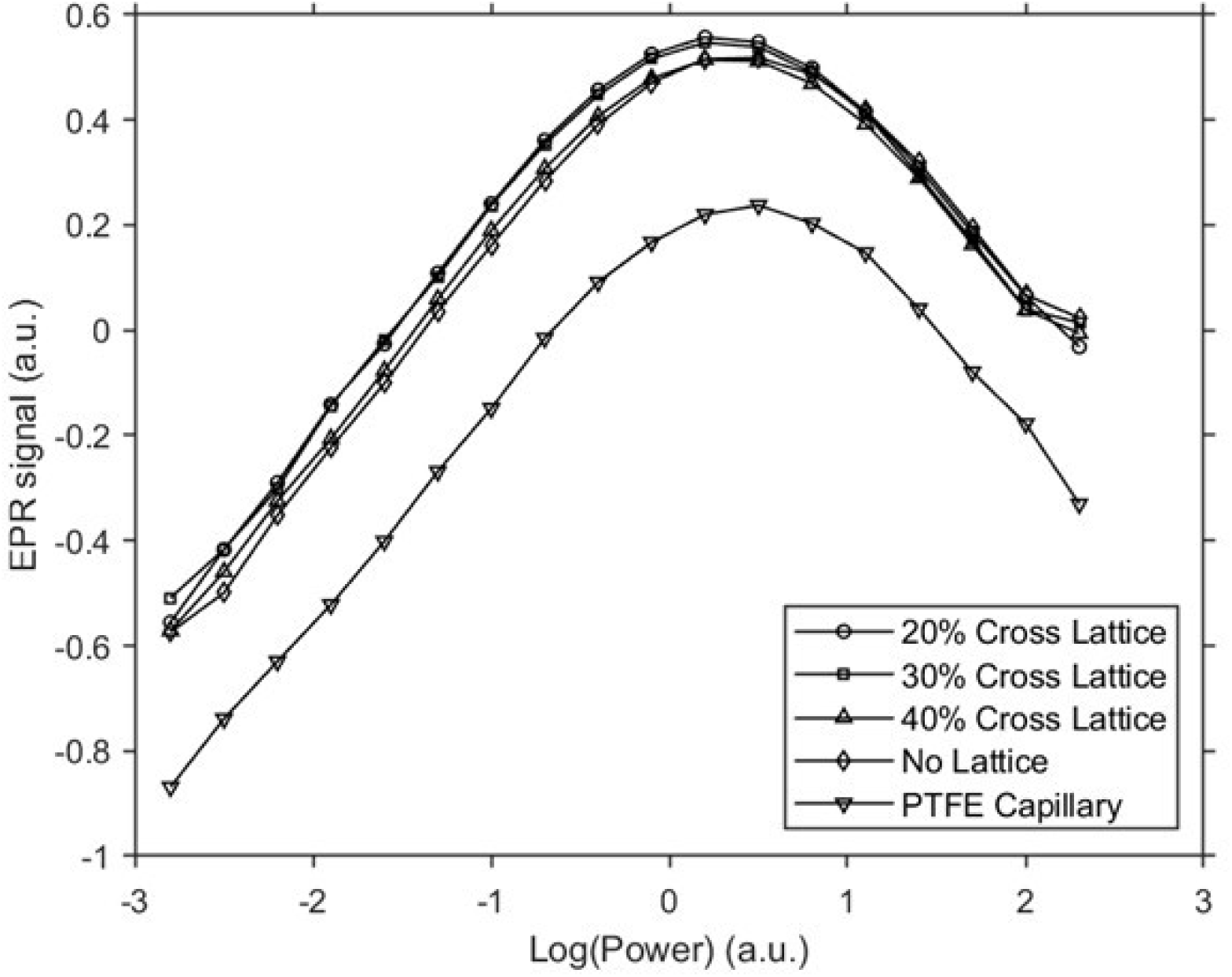
Experimental results of power saturation measurements for cross-type lattice sample tubes. Log(Peak to peak signal) plotted against Log(power), Each measurement was repeated (n = 2) for reproducibility.

**Figure 16:**
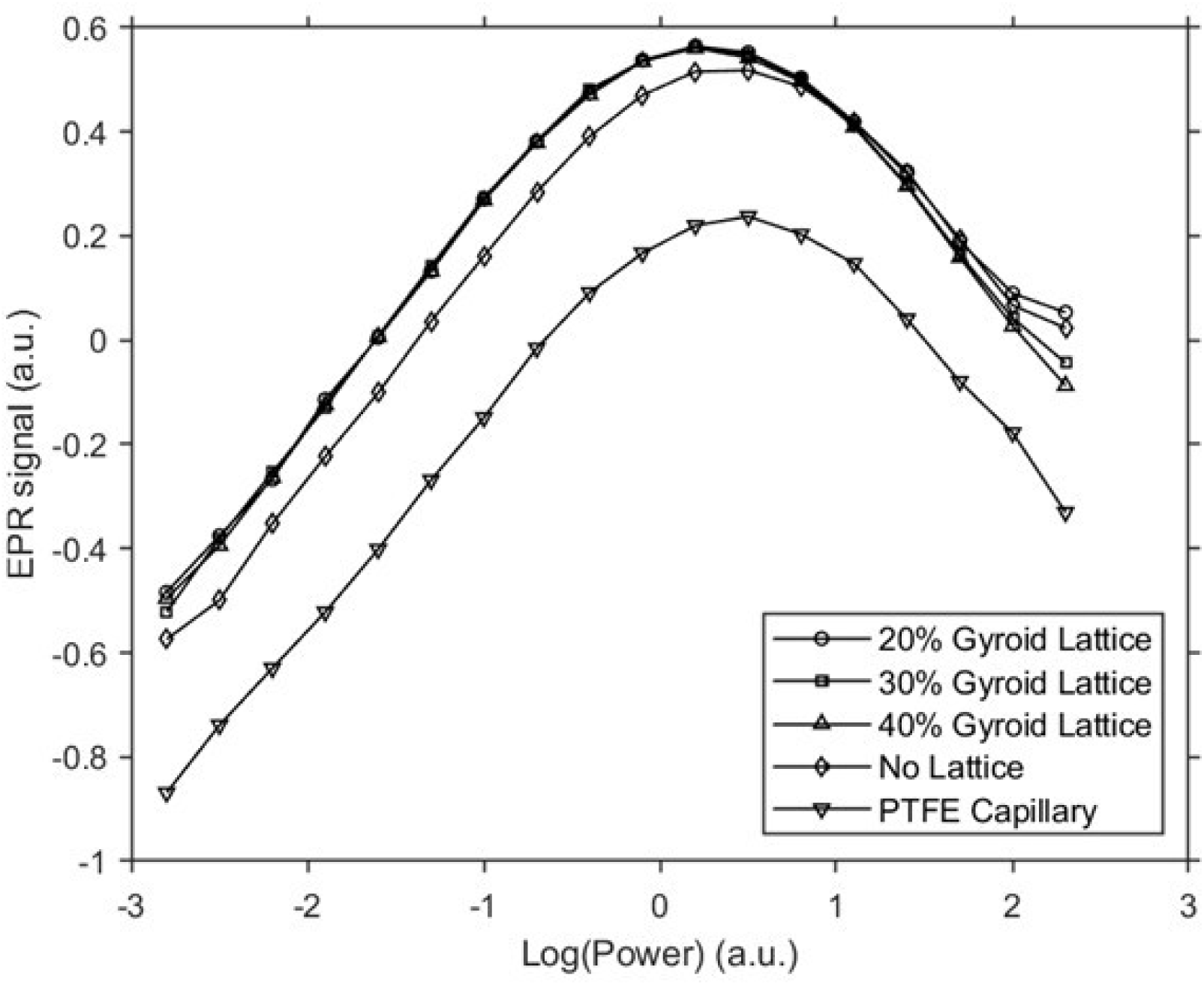
Experimental results of power saturation measurements for gyroid-type lattice sample tubes. Log(Peak to peak signal) plotted against Log(power), Each measurement was repeated (n = 2) for reproducibility.

**Figure 17:**
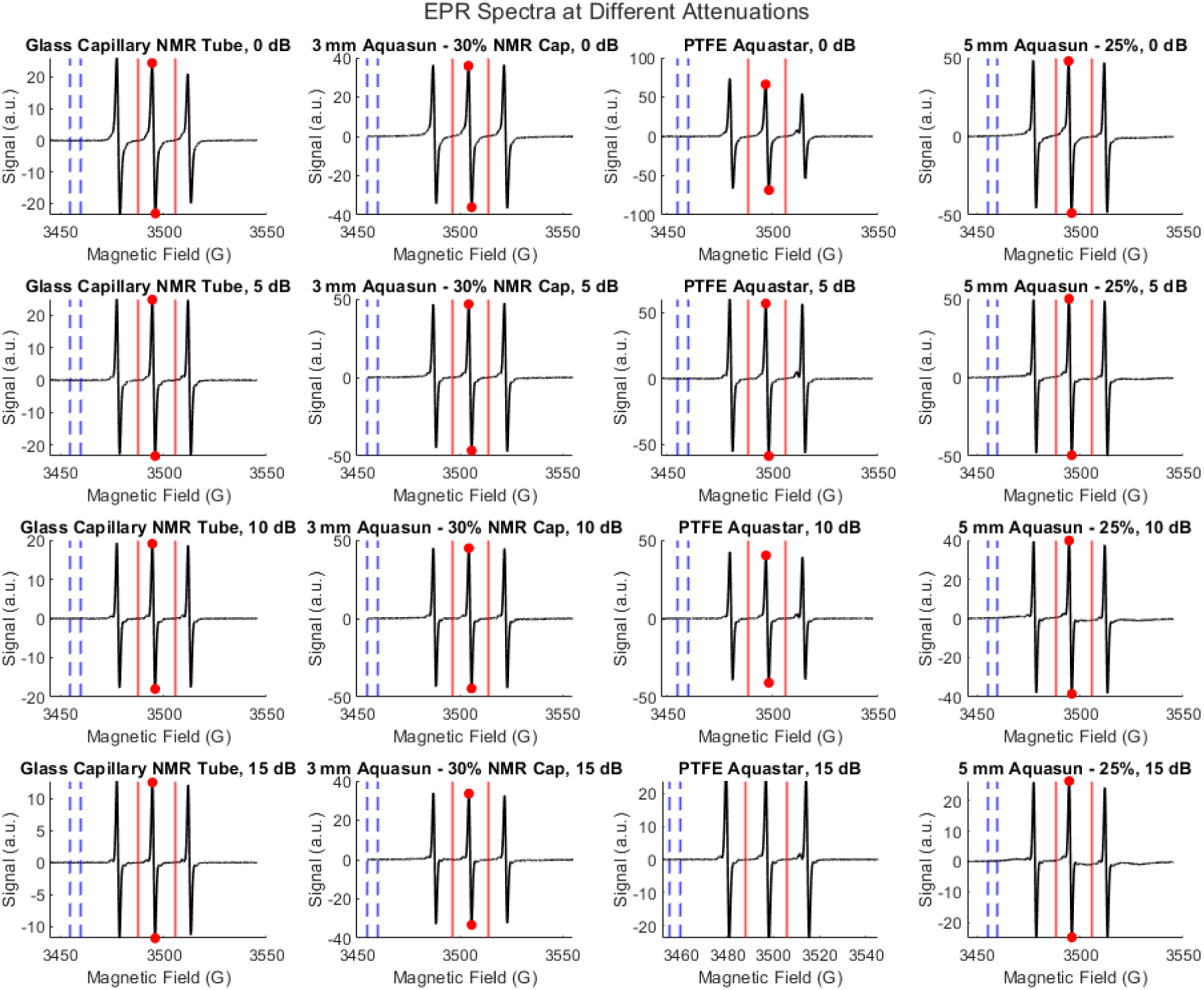
Representative CWEPR spectra of aqueous TEMPOL solution loaded in different sample tubes and at different microwave attenuation. The interval inside the blue dashed line represent the field interval used for SNR calculation. and the red lines represent the field interval used for peak to peak intensity calculation.

**Figure 18:**
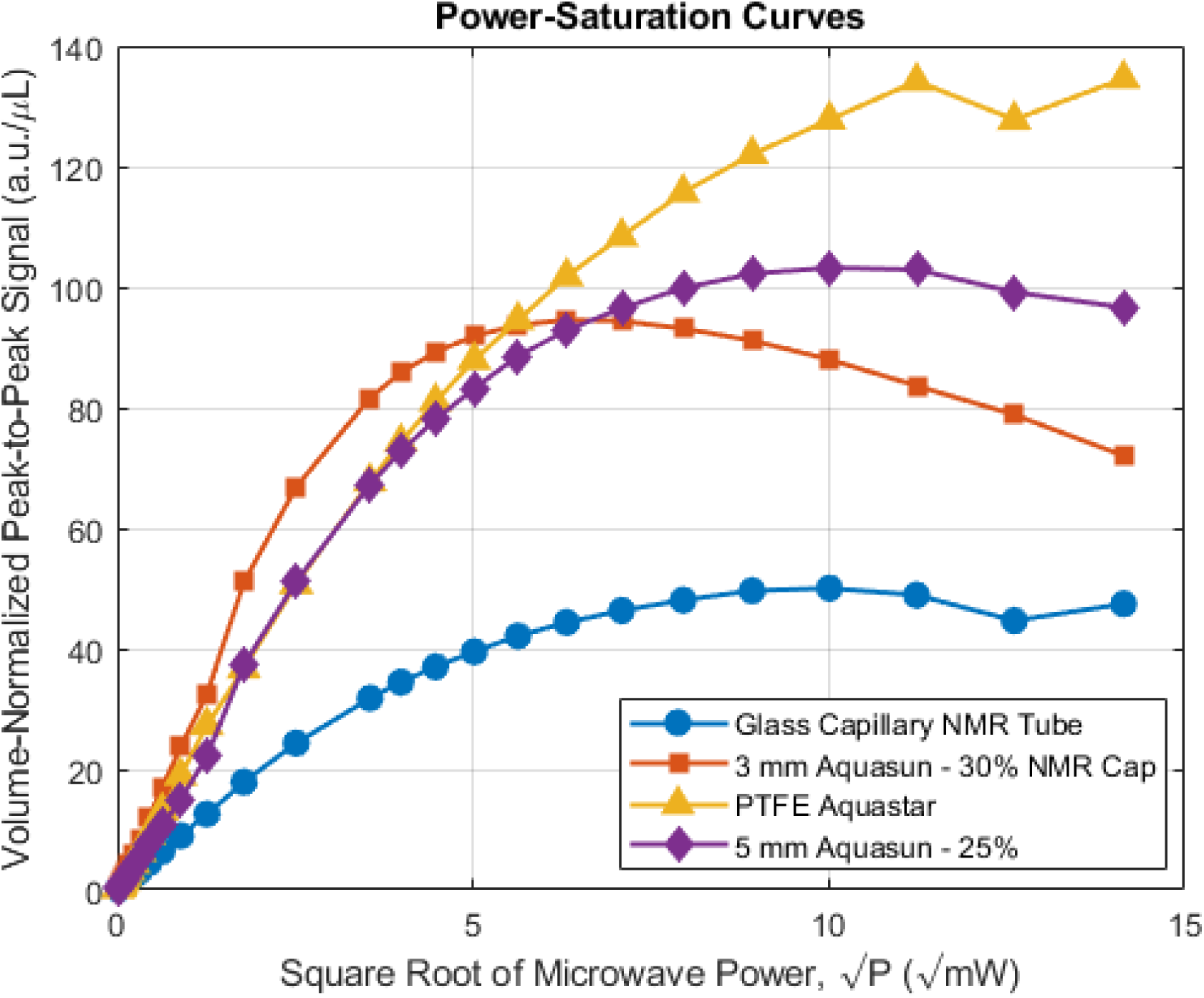
Power Saturation results for aquasun sample tubes using TEMPOL solution.

**Figure 19:**
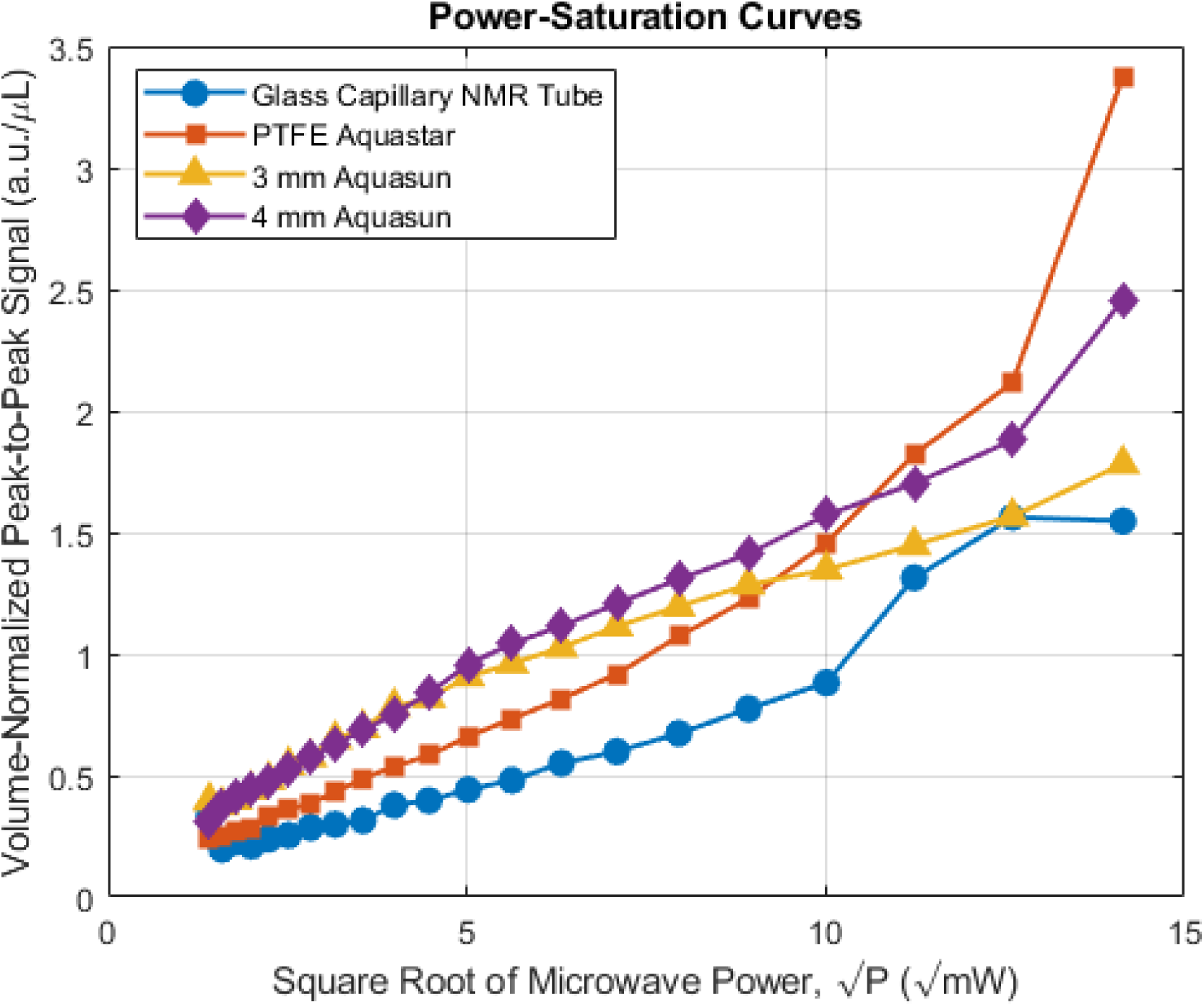
Power Saturation results for aquasun sample tubes using T4L118R1 solution.

**Figure 20:**
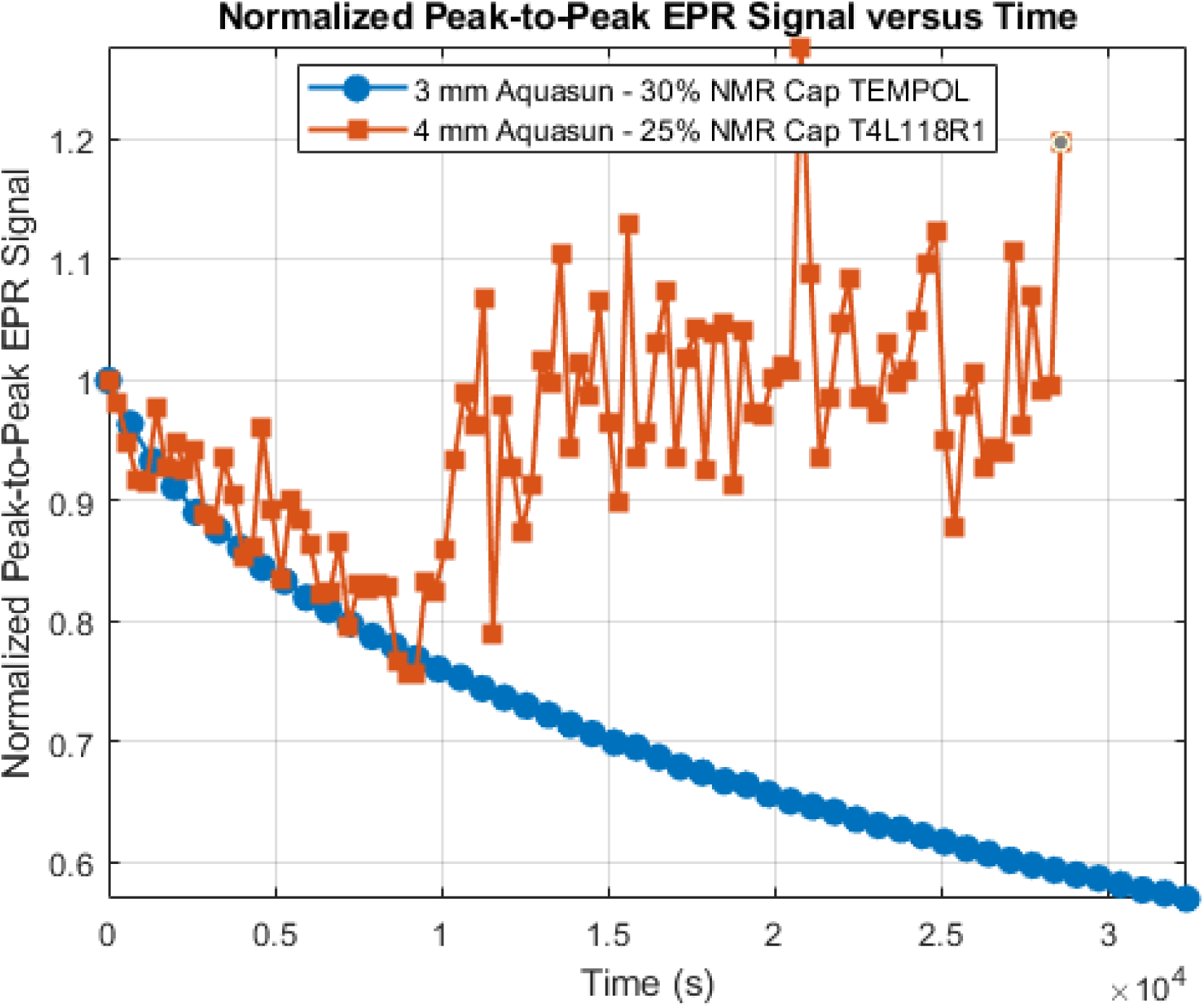
Power Saturation results for aquasun sample tubes using T4L118R1 solution.

**Figure 21:**
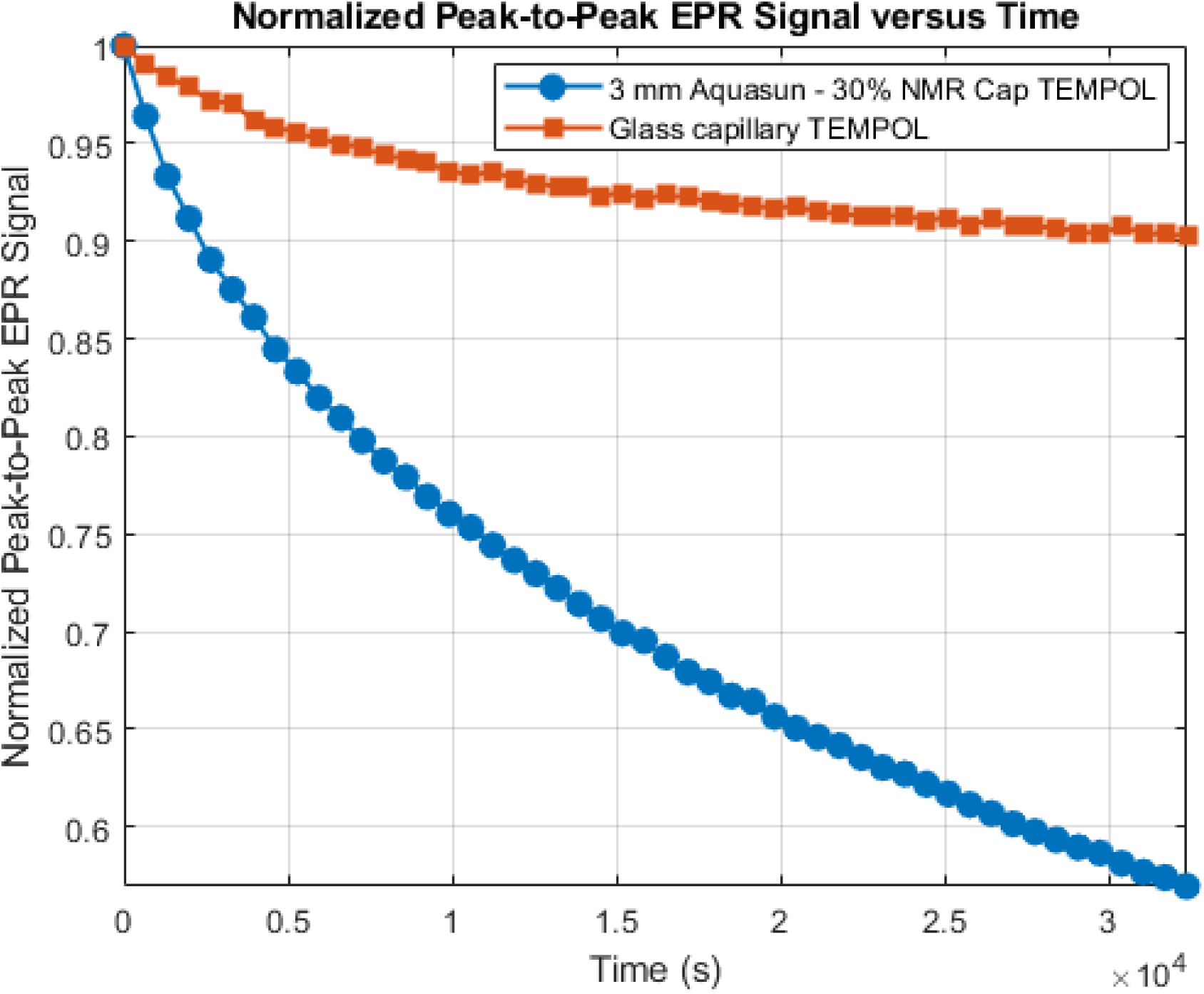
Power Saturation results for aquasun sample tubes using T4L118R1 solution.

## Notes

### Competing Interest Statement

The authors have declared no competing interest.

## References

1. J. M. Hirshon, G. K. Fraenkel, Recording high-sensitivity paramagnetic resonance spectrometer, Review of Scientific Instruments 26 (1) (1955) 34–41. arXiv:https://pubs.aip.org/aip/rsi/article-pdf/26/1/34/19108157/341online.pdf, *doi*: 10.1063/1.1771235*. URL*

[2] R. R. Mett, J. S. Hyde, Aqueous flat cells perpendicular to the electric field for use in electron paramagnetic resonance spectroscopy, Journal of Magnetic Resonance 165 (1) (2003) 137–152. 10.1016/S1090-7807(03)00275-1. URL https://www.sciencedirect.com/science/article/pii/S1090780703002751

3. J. S. Hyde, A new principle for aqueous sample cells for epr, Review of Scientific Instruments 43 (4) (1972) 629–631. arXiv:https://pubs.aip.org/aip/rsi/article-pdf/43/4/629/19018817/6291online.pdf, doi: 10.1063/1.1685709. URL

[4] J. W. Sidabras, R. R. Mett, J. S. Hyde, Extruded dielectric sample tubes of complex cross section for epr signal enhancement of aqueous samples, Journal of Magnetic Resonance 277 (2017) 45–51. 10.1016/j.jmr.2017.02.009. URL https://www.sciencedirect.com/science/article/pii/S109078071730037X

5. S. Misra, Multifrequency electron paramagnetic resonance: Theory and applications (2011). URL https://books.google.com/books?id=VUOnj5s5WAQC

[6] BMF Materials, Htl: Material data sheet (2024).

7. R. R. Mett, A. M. Garces, A. Anilkumar, J. T. Wehrley, M. T. Lerch, C. S. Klug, J. W. Sidabras, Improved epr sensitivity for aqueous biological samples using low-volume multi-channel cells and dielectric resonators, bioRxiv (2025). arXiv:https://www.biorxiv.org/content/early/2025/06/07/2025.06.03.657726.full.pdf, doi:10.1101/2025.06.03.657726. URL https://www.biorxiv.org/content/early/2025/06/07/2025.06.03.657726

8. D. A. G. Bruggeman, Berechnung verschiedener physikalischer konstanten von heterogenen substanzen. i. dielektrizitätskonstanten und leitfähigkeiten der mischkörper aus isotropen substanzen, Annalen der Physik 416 (7) (1935) 636–664. arXiv:https://onlinelibrary.wiley.com/doi/pdf/10.1002/andp.19354160705, 10.1002/andp.19354160705. URL https://onlinelibrary.wiley.com/doi/abs/10.1002/andp.19354160705

[9] V. A. Markel, Introduction to the maxwell garnett approximation: tutorial, J. Opt. Soc. Am. A 33 (7) (2016) 1244–1256. doi:10.1364/JOSAA.33.001244. URL https://opg.optica.org/josaa/abstract.cfm?URI=josaa-33-7-1244

[10] A. Sihvola, Mixing rules with complex dielectric coefficients, Subsurface Sensing Technologies and Applications 1 (4) (2000) 393–415. doi:10.1023/A:1026511515005. URL

[11] Y. E. Nesmelov, A. Gopinath, D. D. Thomas, Aqueous sample in an epr cavity: sensitivity considerations, Journal of Magnetic Resonance 167 (1) (2004) 138–146. 10.1016/j.jmr.2003.12.005. URL https://www.sciencedirect.com/science/article/pii/S1090780703004348

[12] R. R. Mett, J. W. Sidabras, J. R. Anderson, C. S. Klug, J. S. Hyde, Rutile dielectric loop-gap resonator for x-band epr spectroscopy of small aqueous samples, Journal of Magnetic Resonance 307 (2019) 106585. 10.1016/j.jmr.2019.106585. URL https://www.sciencedirect.com/science/article/pii/S1090780719302241

[13] C. Altenbach, C. J. López, K. Hideg, W. L. Hubbell, Chapter three - exploring structure, dynamics, and topology of nitroxide spin-labeled proteins using continuous-wave electron paramagnetic resonance spectroscopy 564 (2015) 59–100. 10.1016/bs.mie.2015.08.006. URL https://www.sciencedirect.com/science/article/pii/S0076687915004589

